# In-vivo fast non-linear microscopy reveals impairment of fast axonal transport induced by molecular motor imbalances in the brain of zebrafish larvae

**DOI:** 10.1101/2022.07.09.499436

**Authors:** Baptiste Grimaud, Maxence Frétaud, Feriel Terras, Antoine Bénassy, Karine Duroure, Valérie Bercier, Gaëlle Trippé-Allard, Rabei Mohammedi, Thierry Gacoin, Filippo Del Bene, François Marquier, Christelle Langevin, François Treussart

**Author notes:** These authors equally contributed to the work (Baptiste Grimaud, Maxence Frétaud, Feriel Terras).

## Abstract

Cargo transport by molecular motors along microtubules is essential for the function of eucaryotic cells, in particular neurons in which axonal transport defects constitute the early pathological features of neurodegenerative diseases. Mainly studied in motor and sensory neurons, axonal transport is still difficult to characterize in neurons of the brain in absence of appropriate *in vivo* tools. Here, we measured fast axonal transport by tracing the second harmonic generation (SHG) signal of potassium titanyl phosphate (KTP) nanocrystals endocytosed by brain neurons of zebrafish (Zf) larvae. Thanks to the optical translucency of Zf larvae and of the perfect photostability of nanoKTP SHG, we achieved a high scanning speed of 20 frames (of ≈ 90 *μ*m×60 *μ*m size) per second in Zf brain. We focused our study on endolysosomal vesicle transport in axons of known polarization, separately analyzing kinesin and dynein motor-driven displacements. To validate our assay, we used either loss-of-function mutations of dynein or kinesin 1 or the dynein inhibitor dynapyrazole, and quantified several transport parameters. We successfully demonstrated that dynapyrazole reduces nanoKTP mobile fraction and retrograde run length consistently, while the retrograde run length increased in kinesin 1 mutants. Taking advantage of nanoKTP SHG directional emission, we also quantified fluctuations of vesicle orientation. Thus, by combining endocytosis of nanocrystals having non-linear response, fast two-photon microscopy, and high-throughput analysis, we are able to finely monitor fast axonal transport *in vivo* in the brain of a vertebrate, and reveal subtle axonal transport alterations. The high spatiotemporal resolution achieved in our model may be relevant to precisely investigate axonal transport impairment associated to disease models.

---

Active intracellular transport of cargoes such as organelles, vesicles, RNA granules and other materials, along the cytoskeleton, is an essential molecular process participating in cell homeostasis. This transport is critical in axons and dendrites of neurons, that form extensive branches connecting different organs of the nervous system. Indeed, axonal transport deficits are hallmarks of neurodegenerative diseases^1^ and recent studies have also consolidated the key role played by molecular transport in the maintenance and functions of synapses.^2^

Axonal transport is categorized as fast or slow at velocities ranging from a few *μ*m/s for organelles to < 0.1 *μ*m/s for cytoskeletal proteins,^3^ respectively. This process takes place along the microtubule (MT) cytoskeleton and is sustained by two types of molecular motors: the superfamily of kinesin motors^4^ ensuring transport of cargoes from MT minus end to the plus end, and the dynein complex^5^ traveling in the opposite direction. As in axons, MT are uniformly polarized with all their plus ends distal from neuron soma, kinesin is responsible for anterograde motions (from soma to axon periphery), and dynein for retrograde displacements (from axon periphery to soma). Over the past few decades, much progress has been made in developing tools and methods to visualize and measure axonal transport.^6^ The most common measurement approach relies on videomicroscopy recording of fluorescently-labeled cargoes displacements. Depending on the type of cargo and the complexity of the biological system (neuronal cell culture, primary neurons *ex vivo* tissue cultures or *in vivo* models), various labeling methods have been selected. The latter include use of live cell compatible dyes like MitoTracker or LysoTracker for mitochondria or lysosomes respectively, fluorescently labeled neurotrophic toxins,^7^ or antibody against neurotrophin^8^ or fluorescent viral particles.^6,9^ Furthermore, taking advantage of genome editing tools applied to model organisms like *Drosophila* and *Danio rerio* (Zebrafish, Zf), fluorescent proteins (FP) have been expressed in specific organelles of given neuronal populations to investigate their axonal transport,^6,10,11^ in different contexts including neurodegenerative diseases,^12,13^ axonal regeneration^14^ and age-dependent decline. ^15^ These physiological models, particularly relevant for the study of complex biological processes by imaging, can be delicate to implement. First of all they rely on transgenesis by injection of the transgene constructs at the single cell level to obtain the mosaic expression^11–13^ of the marker of interest, necessary to allow single organelle identification. Then, injected larvae must be selected based on the expression pattern of the marker in terms of localization and level, to be used for axonal transport investigation.

Finally, more challenging studies were also conducted *in vivo* in transgenic mice to monitor the axonal transport of fluorescent mitochondria in peripheral nerves, ^16,17^ but they required a complex and invasive surgical exposure of the nerves, animal immobilization and optimization of fast and 3D deep imaging processes.^10^ Axonal transport was also measured in neurons of mouse central nervous system (CNS) using two-photon excited microscopy. This includes neurons of the spinal cord,^18^ retinal ganglion cells^19^ and cortical neurons.^20^ However, these approaches not only still require some surgery but photobleaching of the FP reporter and phototoxicity for the animal also limit the temporal resolutions to a maximum of one frame per second, with a spatial resolution of about 200 nm. This time resolution is sufficient to evaluate parameters such as the fraction of mobile cargoes, their velocity and direction of transport but it prevents the observation of short pauses (duration ≲ 250 ms) which are of utmost interest to investigate the microtubule (MT) network complexity and stability,^3^ that were recently reported by Chowdary *et al*.^21^ and by us in endolysosomal transport in cultured neurons, using a novel nanoparticle tracing methodology^22^ acquiring at 20 frame/s (fps). These micro-pauses were attributed to tug-of-war situations between kinesin and dynein, both attached to the same cargo and pulling in opposite direction, leading occasionally to short pauses when the cargo experiences null net force. ^23^ The present study aims at investigating accurately axonal transport modifications of endolysosome compartments induced by modulations of molecular motors concentrations in zebrafish larvae brain, *in vivo*. To be sensitive to short pauses, it is therefore critical to achieve a frame rate at least equal to the 20 fps used in sensitive *in vitro* studies. ^21,22^ Indeed intravital imaging of organelle transport can also benefit from the use optically-active photostable nanocrystals internalized in neuronal cells by endocytosis and subsequently traced at large frame rate. This technology was pioneered by Cui *et al* ^24^ who used quantum dot as fluorescent tracers. Later, we reported the use of fluorescent diamond nanocrystals as tracers to reveal subtle changes in endolysosomal transport within cultured neurons established from a transgenic mouse bearing a genetic risk factor of a neuropsychiatric disease. ^22^

Here we extended such nanoparticle-based assay to measure the endolysosomal transport parameters *in vivo*, in neurons of the brain of intact, living zebrafish larvae. To this aim we used potassium titanyl phosphate (KTiOPO_4_, KTP) nanocrystals (size ≈ 120 nm) that exhibit large second-order non-linear optical response, and micro-inject them in optic tectum of Zf larvae. Incidentally, the ability to image these efficient nonlinear KTP nanocrystals (nanoKTP) in live Zf larvae blood circulation at ultra-high frame rate has been recently reported in a wide-field configuration with light-sheet illumination. ^25^ Furthermore, in a previous work, we had shown that nanoKTP are spontaneously internalized in 2D primary cultures of neurons and can be traced by acquisition of their second harmonic generation (SHG) emission under infrared (IR) pulsed laser excitation.^26^ Key advantages of SHG over fluorescence are non-saturation, non-bleaching, and a reduced background, hence a large signal to background ratio. In addition, the wavelength of SHG emission is tunable and can be adjusted to avoid any overlap with other fluorescent labels used in the same sample.

In the present work, we harnessed these properties combined with fast raster scanning of the IR laser beam to achieve the frame rate of 20 frames/s (fps) for a ≈ 80 *μ*m-sized field-of-view, identical to the one employed in our *in vitro* intraneuronal transport assay. ^22^ Moreover, due to a single dominant coefficient of the nonlinear susceptibility tensor, ^27^ SHG from KTP has a directional emission. We took advantage of this property to investigate the vesicle rotational dynamics during their axonal transport, in a similar way as it was reported in cultured neurons, ^28,29^ where very fast dynamics was captured at 500 fps (2 ms frame duration), justifying further our settings to the highest possible frame rate. The amplitude of vesicle rotational motions is an indirect readout of how strongly kinesin and dynein motors tether it to microtubules. The more motors there are, tethering a vesicle to a microtubule, the more constrained are its position and its orientation relative to a reference frame, as the motor counteract Brownian motion. If some motors linked to the vesicle untether from the microtubule, the amplitude of vesicular Brownian motions, that include rotations, will increase. We hypothesize, like in Kaplan et al.,^29^ that the amplitude of orientational fluctuations is a qualitative readout of the number of motors attached to both the endosome and the microtubule, and therefore that it adds significant value to the analysis, at no additional experimental cost.

In our experiment, we showed that the nanoKTP move within axons of periventricular neurons (PVN) that all project radially inside the neuropil to connect retinal ganglion cells.^30^ Considering this anatomical characteristics of PVN, we were able to distinguish the anterograde transport, from the retrograde transport, and to quantify associated parameters separately for each direction. While in our previous study in cultured neurons, we had investigated the impact on intraneuronal transport of microtubule disruption or of the level of a kinase phosphorylating MT-associated proteins, without distinction of axons and dendrites, here we focused on active molecular motor-dependent mechanisms. As we addressed specifically axonal transport along polarized MT, we could relate the anterograde or retrograde transport direction to the activity of one type of motors. We considered transgenic Zf engineered to bear loss-of-function alleles of retrograde motor protein Dync1h1 heavy chain 1 of the dynein motor complex^31^ or of the anterograde motor protein Kif5aa.^32^ As in *dync1h1* mutant all transport was halted, we opted for a drug inhibition strategy to achieve a finer control of retrograde transport, using dynapyrazole, a cell permeable, conformationally constrained isostere of ciliobrevin recently described *in vitro* to perturb retrograde transport.^33^

The comprehensive measurements of translational and rotational motions in these models provided by our nonlinear nanoparticle-based assay opens prospects to study finely axonal transport in the central nervous system *in vivo*. In particular our method may be instrumental to investigate the biological significance of altered cargo pausing duration and run length and to study the link between transport defects and neuronal dysfunctions. In this context, the zebrafish model will be amenable to conduct high-content screening of therapeutic drugs able to rescue axonal transport impairment.

## Results and Discussion

### *In vivo* measurement of intraneuronal transport

To investigate the axonal transport in the CNS of Zf larvae imaged by fast non-linear microscopy, we synthesized single crystal nanoKTP (Supporting Figure S1A) in solution in deionised water as we reported previously. ^26^ Using differential centrifugation we narrowed their size distribution, that was measured by nanoparticle tracking analysis, leading to an average size of 123.8± 2.4 nm, with a standard deviation of 42.7 nm (Supporting Data S1 and Figure S1B). This solution was injected in the left optic tectum of 3 days post-fertilization (dpf) larvae at a concentration of ≈ 1 mg/mL (Fig. 1A). We conducted the non-linear microscopy fast-scanning recording 24 hours later, at 4 dpf, when tectal neurons are mature and respond to visual stimulations. ^34^ Prior to live imaging of the optic tectum, Zf larvae were anesthetized and placed in dorsal view in an agar mold (Fig. 1B). Once in the good orientation, larvae were immobilized by covering them with a cushion of low-melting agarose maintained during the acquisition, performed at 24°C. The injection condition (solution concentration and volume) were optimized to obtain a sparse labeling, but sufficiently dense to collect enough data to ensure reliable statistical analysis in order to minimize the risk of interference with the endocytosis along the axons, which is a key physiological process for neuronal polarity maintenance. ^35^ Failure of the injection was rare: about 90% of injected larvae contained enough nanoKTP in the region of interest for axonal transport analysis.

**Figure 1:**
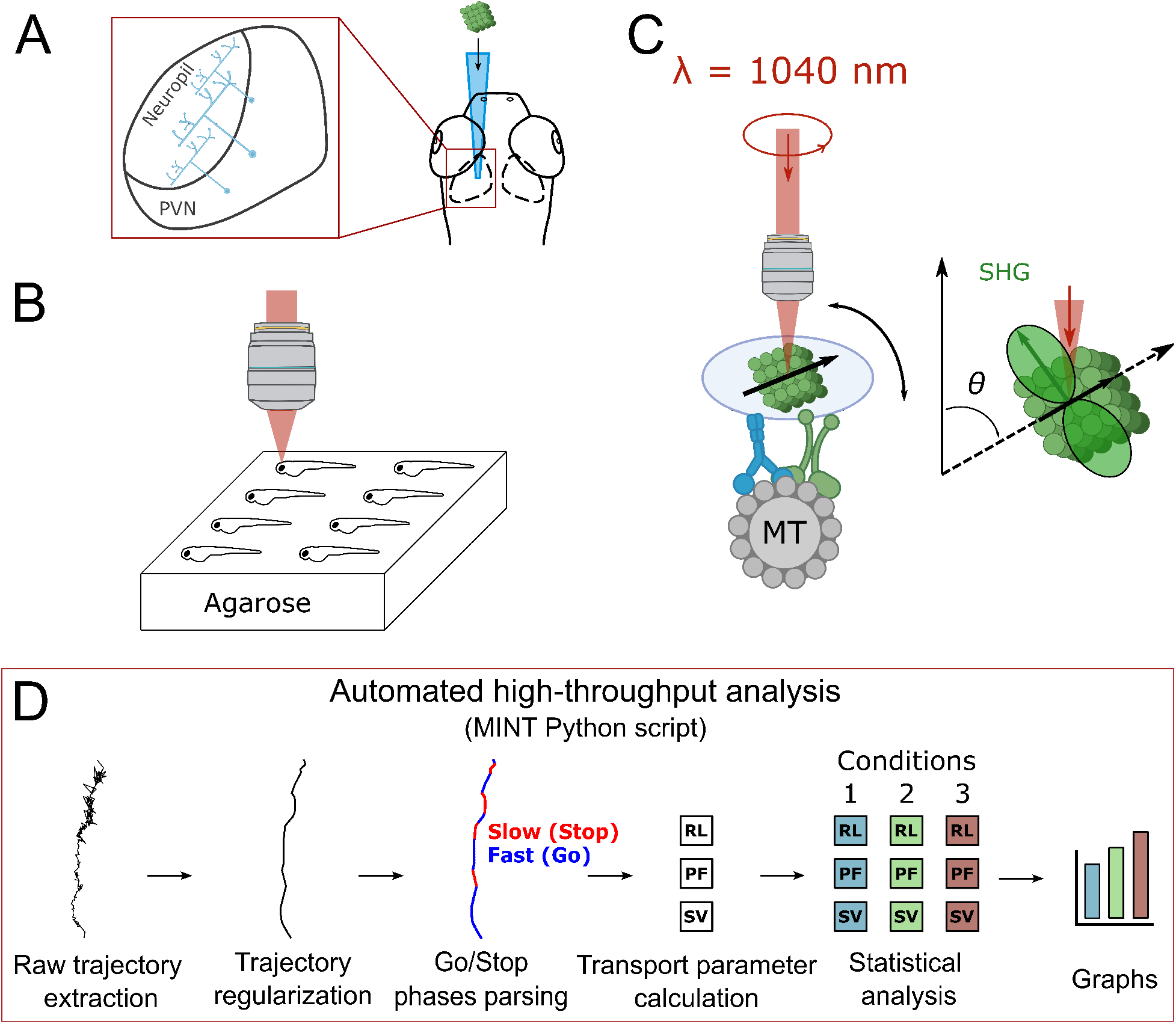
Method to measure intraneuronal transport parameters in the brain of zebrafish larvae, based on non-linear nanocrystals microinjection and endocytosis, second-harmonic generation fast microscopy, and high-throughput automatic extraction and analysis of nanocrystal-labeled vesicle trajectories. (A) Microinjection of the nanoKTP (green cube) in the left optic tectum of the head of a zebrafish larva (at 3 dpf). Inset: zoom on optic tectum displaying periventricular neurons (PVN) projecting parallel and coplanar axons in the neuropil. (B) Immobilization at 4 dpf of 8 anesthetized injected larvae in channels molded in agarose, with dorsal orientation towards the microscope objective for SHG microscopy. (C) Left: schematics of a single monocrystalline nanoKTP embedded in a vesicle connected to a microtubule (displayed as a section, MT) by a dynein (blue) and a kinesin (green). The black arrow onto the nanoKTP indicates the direction of the second order non-linear polarization (mostly dipolar), in response to the pulsed-laser excitation at 1040 nm wavelength. Right: in green, the SHG radiating pattern of one nanoKTP. (D) Analysis pipeline with the Python program MINT we developed,^36^ consisting, from left to right, in two parts of analysis. The first part includes (i) trajectories extraction, (ii) trajectory regularization (filtering out nanoparticle position noise) and (iii) parsing in fast (“Go”) and slow/pause (“Stop”) phases of motion. The second part starts with (i) the calculation of transport parameters for each trajectory, among which the segmental velocity (SV), which is the average velocity during a Go phase segment, the run length (RL), which is the distance traveled between two consecutive Stops, the pausing frequency (PF), etc and continuing with (ii) the analysis of the data for different biological conditions provided as inputs, transport parameters statistical comparison, and finally (iii) the drawing of graphs associated to these data.

We set the excitation laser wavelength at 1040 nm, and its power at 12.5 mW at the exit of the microscope objective (×25, water immersion, numerical aperture 0.95). This power value was chosen to take advantage of the full dynamic range of the detector (8-bits encoding, maximum pixel intensity of 255). As the nanoKTP SHG is mainly excited by an electric-field direction along a single crystalline axis we use a circularly polarized excitation beam (Fig. 1C) in order to favor the excitation of all the particles whatever their orientation. The light from the nanoKTP is collected with the same microscope objective, then passed through a bandpass filter (50 nm spectral width) centered on 525 nm wavelength, before being detected by a point detector, without any polarization selection. Taking advantage of prism-based tunable emission spectrum detection of our microscope, we recorded the signal emission spectrum from a single nanoKTP and confirmed that it corresponds solely to SHG signal (Supporting Data S2 and Figure S2).

Driving the mirror galvanometers of the microscope at their resonance frequency (8 kHz), we acquired continuous 512 × 352 pixels raster scans at a constant depth position (≈ 100 *μ*m) below the skin with a line average of 2, leading to a refreshing rate of 20 frames/s (*i.e*. 50 ms per full frame scan) identical to the one we use in wide-field acquisition on cultured neurons,^22,37^ that enhances the sensitivity to short pauses. In order to avoid degradation of the diffraction limited spatial resolution by sampling, we selected a pixel size of 173 nm, which fulfills the Shannon criteria as it is smaller than half the diffraction-limited spot radius of two-photon excited microscopy 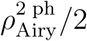. The Airy radius in two-photon excited process can indeed be estimated by 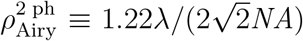, leading to 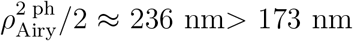, at the excitation wavelength of *λ* = 1040 nm and for *NA* = 0.95 microscope objective numerical aperture.

With this pixel size value of 173 nm, the scan covers 88.6 *μ*m × 60.9 *μ*m. We set the total number of frames in the video to either 2354 (Supporting Video S1) or 2501 (Supporting Video S2), which corresponds to video durations of about 120 s. Compared to the wide-field mode, for which the integration time per pixel is 50 ms, here in the raster-scan mode, the duration of photon counting (dwell time) is only 144 ns/pixel (taking into account the 2-lines averaging done while scanning). To collect a sufficient number of trajectories per larvae, we record 4 videos corresponding to different regions of interest (and different depths) in the optic tectum ipsi- or contra-lateral to the injection site (total of 8 videos per larva). We did not observe mortality due to intracerebral injections or imaging. The acquired videos are then processed by an automated high-throughput analysis pipeline (Fig. 1D) detailed later.

### Internalisation and distribution of the nanoKTP in the periventricular neurons of the CNS

Figure 2A displays the maximum projection in time of Supporting Video 1. It shows that nanoKTP motions appear mostly parallel and oriented radially relative to the eye (top left corner), as expected in this brain region (Fig. 1A, inset). This suggests their internalization in neurons, whose projections line the axons of PVN. Here we stress that this endolysosomal vesicle labeling strategy (via nanoKTP spontaneous uptake) is naturally sparse and does not require further sorting.^11–13^ To better identify the cell type that was targeted by the nanoKTP, we performed simultaneous live labeling of nanoKTP with the cell membrane dye DiI. The pattern of expression obtained in two-photon excited fluorescence microscopy reveals DiI positive cells with the morphology of PVN and the presence of nanoKTP in the axon (Fig. 2B). To confirm these observations, we conducted immunofluorescence and quantified the localization of the nanoKTP in different cell subtypes present at the site of injection. Twenty four hours post injection, Zf larvae were immunostained to label acetylated-tubulin (ac-tub), considered as a marker of axons, and glial fibrillary acidic protein (GFAP), as a marker of glial cells and neuronal precursors (Fig. 2C-E). Nanoparticles colocalize in 27% of the cases with ac-tub-positive axons, which is 2.7 times more than the colocalization with GFAP-positive branches presenting 10.2% of colocalization (Fig. 2F). In about 60% of the cases, the nanoKTP were found aside from ac-tub and GFAP-positive structures, which might correspond to particles stuck in the extracellular matrix, unable to diffuse to axonal membranes where they may be endocytosed.

**Figure 2:**
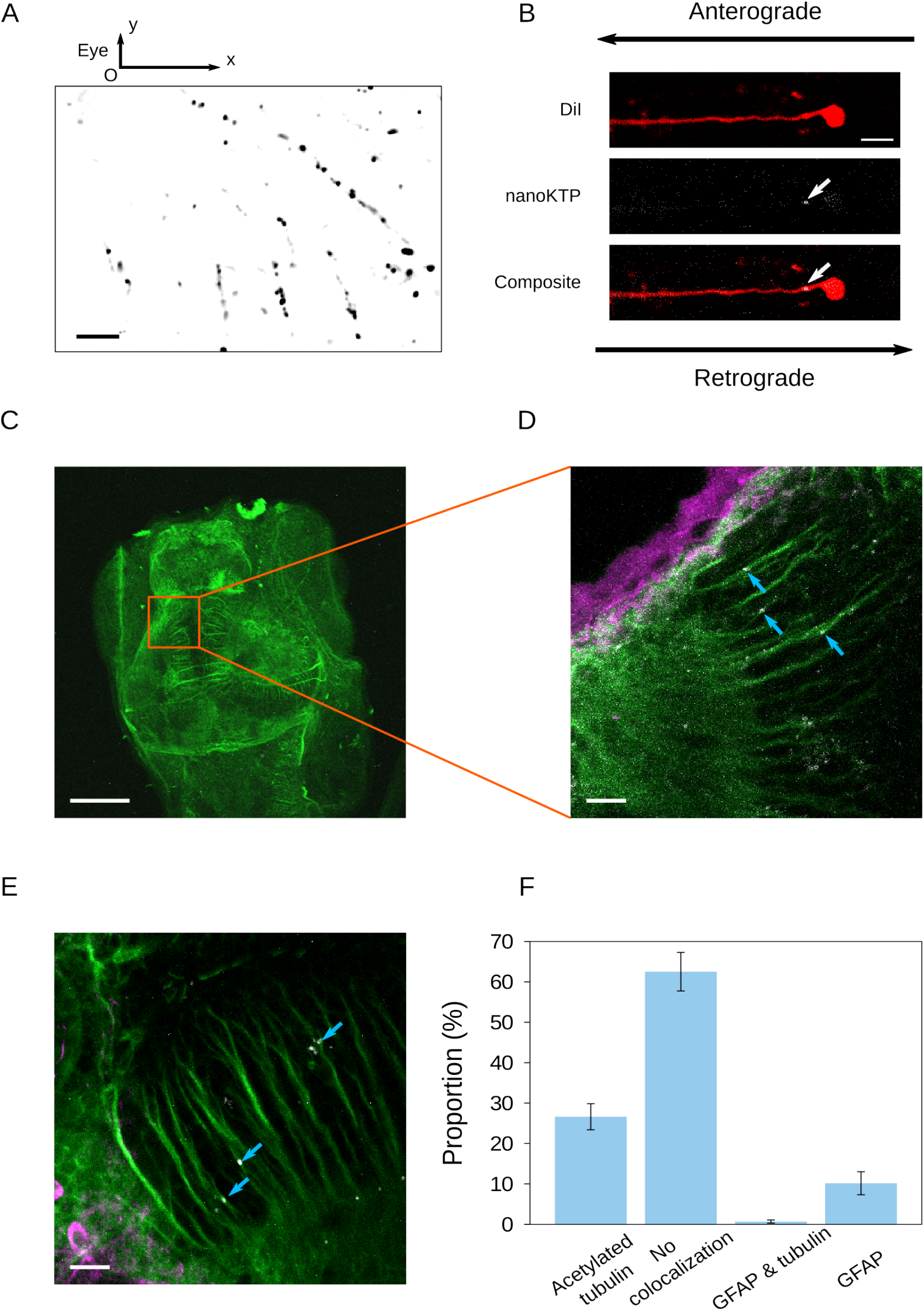
Localization of the nanoKTP relative to axons and glial cells of the zebrafish larva optic tectum injection zone. (A) Average intensity projection of a single plane raster-scan video (Supporting Video 1) of the SHG signal from nanoKTP in the brain of a Zf larvae (at 4 dpf). Moving nanoKTP have directed, mostly linear, displacements oriented towards the left top corner where the larva left eye is located (out of the field of view, FoV). Scale bar: 10 *μ*m. (B) Top: two-photon fluorescence image of a DiI cyanine dye-labeled periventricular neuron in a live Zf (4 dpf); middle: SHG signal of the same FoV as the top panel, showing in white a nanoKTP (white arrow) colocalized with the axon of the PVN. Bottom: merge of top and middle images. Retrograde and anterograde directions are indicated by arrows. Scale bar: 10 *μ*m. (C) Acetylated tubulin immunofluorescence labeling of a zebrafish larva head (in ventral position). Maximum projection of confocal slices of a *z*-stack covering a thickness of 30 *μ*m. Scale bar: 100 *μ*m. (D) Representative field of view (zoom of (C) larva) merging three detection channels: acetylated tubulin (ac-tub, green) and GFAP (magenta) immunostainings, and nanoKTP (white). Maximal projection over a total thickness of 5.1 *μ*m. Nanoparticles colocalized with ac-tub are pointed by arrows. (E) Another FoV with the same labels as in (D). Maximal projection over a total thickness of 4 *μ*m. Scale bars for (D) and (E): 10 *μ*m. (F) Proportion of the localization of nanoKTP relative to ac-tub and GFAP immunostaining (*n* = 4 larvae and 2-7 FoV per larva). Error bars: s.e.m.

As all PVN bodies are gathered in the PVN-labeled area of Fig. 1A inset (between the neuropil and the ventricle) and project towards the neuropil (located above the eye), motions towards negative values of *x* ((O*x*) axis pointing right, with origin in the eye, see Fig. 2A), are anterograde if we are addressing the left eye, and retrograde for the right eye.

Note that in Fig. 2A, most lines are made of a succession of bright and dim spots. This blinking reflects the rotational motion (relative to the microscope objective axis (O*z*)) of the vesicle-containing nanoKTP, upon their transport by the molecular motors stepping on the microtubules. This intensity fluctuation is a consequence of the anisotropy of nanoKTP SHG radiation pattern that can be assimilated to the one of a dipole (Fig. 1C). Indeed KTP second-order nonlinear susceptibility matrix^38^ as a dominant coefficient leading to a dipolar non-linear polarization mostly parallel to the *c* crystallographic axis. When the dipole is aligned with the objective axis, almost no light is collected, leading to a very dim signal. Incidentally, the large variation of intensity of the vast majority of spots indicate that each endosome most probably contain a single nanoKTP, because the presence, within a vesicle, of multiple randomly oriented nanocrystals would yield a much lower intensity contrast. This observation confirms that *in vivo*, like in primary cultures, ^22,37^ neuron internalize at most one solid nanoparticle per vesicle. The nanoKTP still appear as single particles per transported vesicular compartment 24 hours after injection, despite the possibility for endolysosomes to merge. This is probably due to the sparse labeling we use: only a small proportion of vesicles are labeled so it is highly unlikely that two endolysosomes destined for fusion would both contain a nanoKTP.

In the subsequent studies, we evaluated the fraction of moving particles in large sets of data and found 6.7 ± 0.6% (standard error on the mean, s.e.m.) (Fig. 4B) and 8.6 ± 0.8% (Fig. 5B) in two independent studies, leading to an average of 7.7%. We can conclude that, statistically, a non-negligible fraction of 7.7/27 ≈ 28% of the nanoKTP colocalizing with axons, moves during the 2-minutes raster-scans recording. This value must be compared to the mobile fractions of ≈ 60% reported for various endosomes in motor neurons of Zf larvae spinal cord. ^12^ Beyond the fact that our study addresses axonal transport in CNS neurons, that may have different dynamics compared to motor neurons, but also different endolysosomal compartments than in Ref.,^12^ the 2-fold discrepancy observed may be due to an overestimation of nanoKTP internalized in axons. Indeed diffraction limited imaging of sub-micrometer sized structures like axons and nanoKTP makes it difficult to discriminates between particles that stayed at axonal outer-membrane from the ones that were fully internalized.

**Figure 3:**
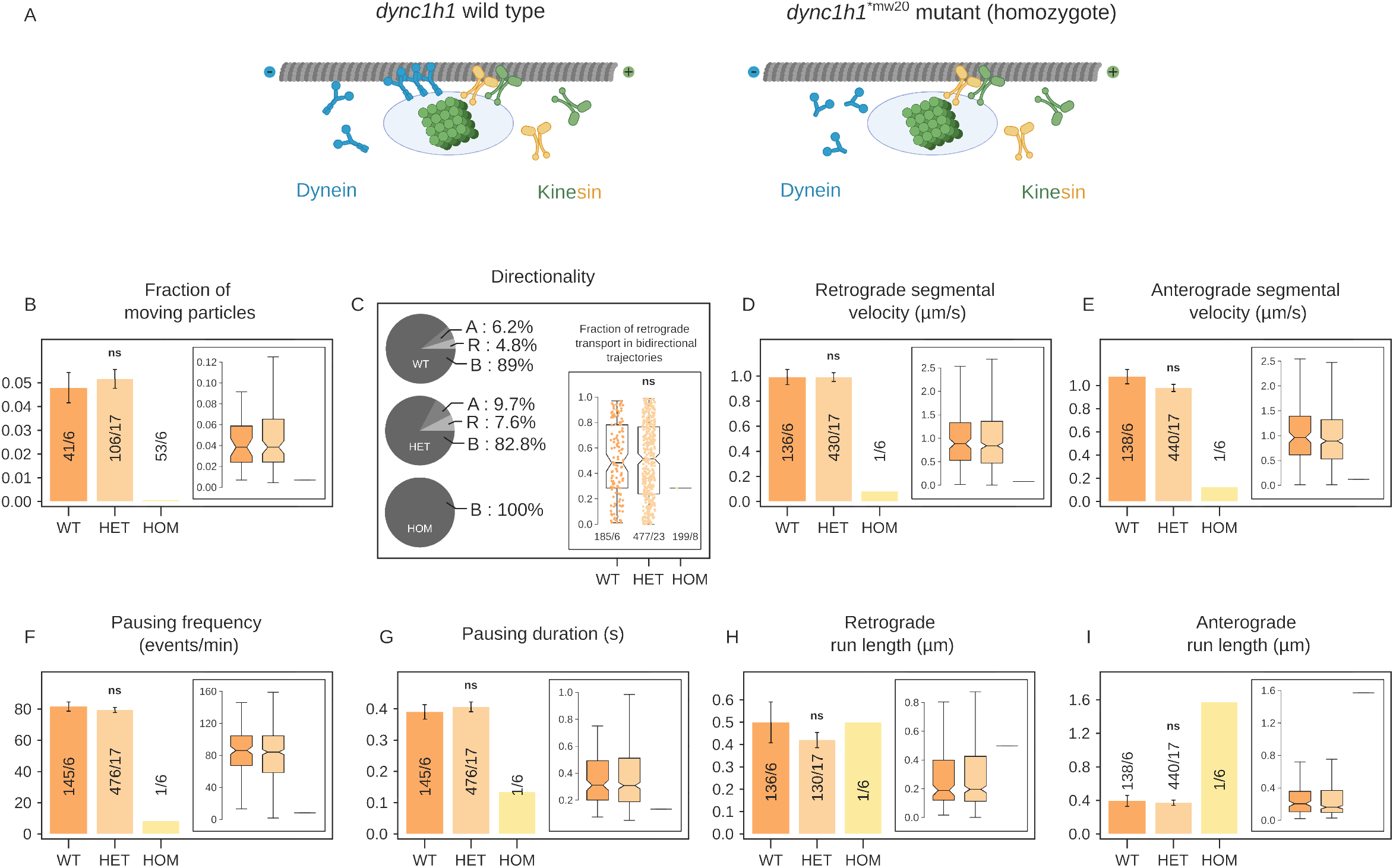
*dync1h1*^mw20^ dynein mutation stops intraneuronal transport in zebrafish brain. (A) Schematic representation of the mechanisms of intraneuronal transport. Left: Typical transport conditions in a wild type *dync1h1*^mw20^ sibling. The optically active nanoparticle (green cube) is spontaneously internalized in an endolysosomal vesicle (light blue oval) and is then transported along a microtubule (grey tube) in an anterograde motion by kinesins (green and yellow) or in a retrograde motion by dyneins (blue). Right: Impaired transport conditions by mutant dynein heavy chain (green shortened dimers). (B–I) Modifications of intraneuronal transport parameters in *dync1h1* ^mw20^ larvae brain, depending on the larva genotype (WT: wild type, HET: heterozygote and HOM: homozygote). For (B, D-I) bar plots display the parameter mean values ±s.e.m., while insets are distribution box plots of the same parameter. In (B) numbers in the bars are # of fields of view analyzed/# of animals analyzed, while for (C-I) they are # of trajectories/# of animals analyzed. (B) Fraction of moving particles, with directed motion. The 3-conditions Kruskal-Wallis *p*-value=0.22. The mobile fraction in HOM mutant is 0.01% while it is 4.8% and 5.2% in WT and HET respectively. (C) Directionality. Left: pie charts of the percentage of purely anterograde [A], purely retrograde [R], or bidirectional trajectories [B], for each condition. Right: fraction of R transport in bidirectional trajectories. No difference are observed (*p* = 0.74). (D-E) Retrograde and anterograde segmental velocities do not vary between WT and HET (*p* = 0.84 and *p* = 0.26). Overall Kruskal-Wallis test *p*-value =0.17 for A and *p*-value =0.29 for R. The A and R phases in the only bidirectional trajectory have very slow segmental velocities of 0.07 *μ*m/s for R and 0.12 *μ*m/s for A. (F) Pausing frequency. Overall Kruskal-Wallis p-value = 0.23. PF does not vary between WT and HET (*p* = 0.49). In HOM the pausing frequency is ten times smaller than in WT and HET (8.2 events/s compared ≈ 80 events/s) (G) Pausing duration does not differs between conditions (*p* = 0.347). There is no difference between WT and HET (*p* = 0.836) but the pausing duration is 3-fold smaller in the trajectory of the HOM mutant. (H) Retrograde run length does not differ between conditions (*p* = 0.572). (I) Anterograde run length does not vary across the different conditions (*p* = 0.21). The RL of ≈ 1.6 *μ*m for the only HOM trajectory is not representative. RL values of ≈ 0.4 *μ*m for WT and HET are consistent with the one reported for *kif5aa* and DYNA data.

**Figure 4:**
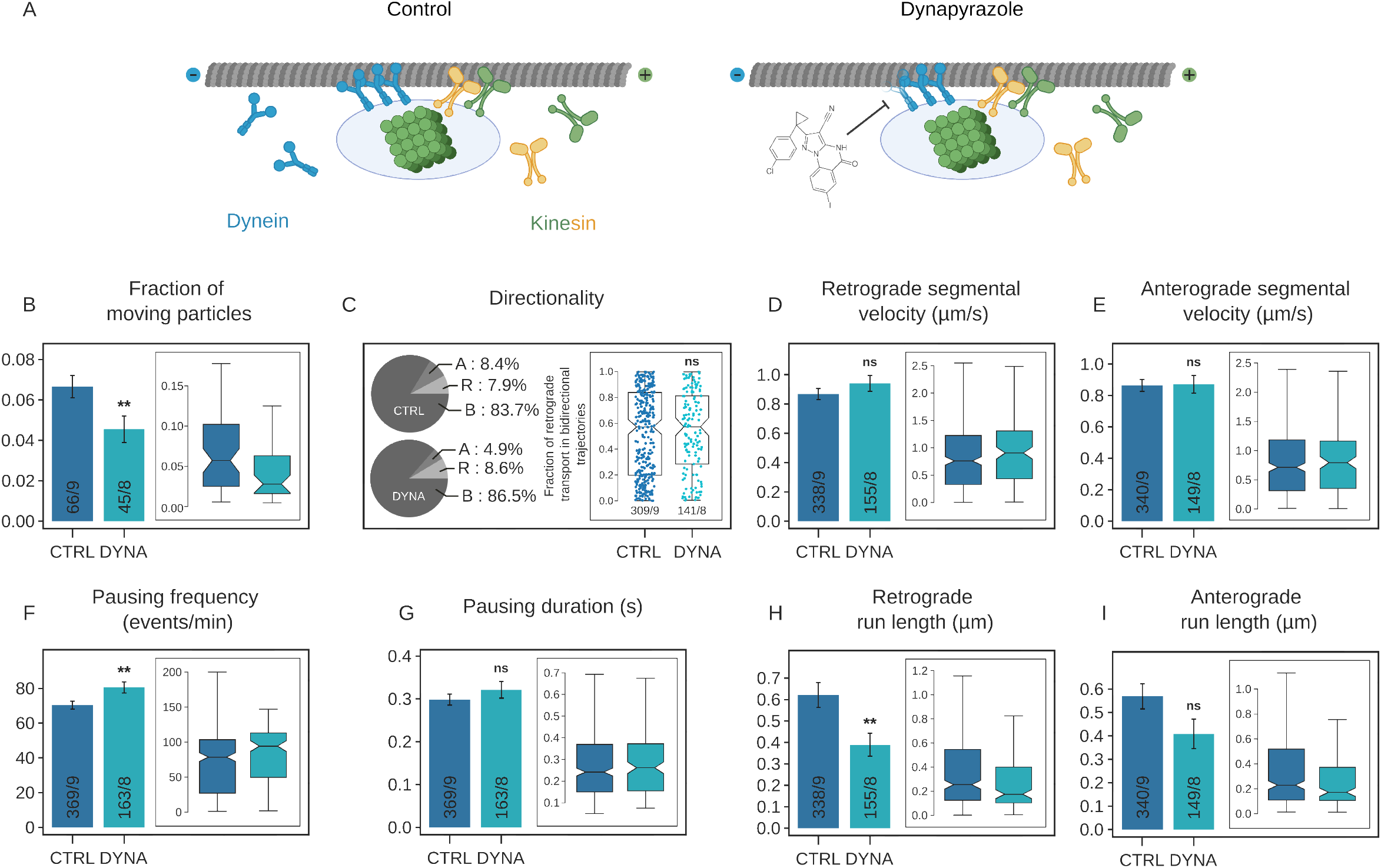
Dynein inhibitor dynapyrazole impairs intraneuronal transport in zebrafish brain. (A) Schematic representation of the mechanisms of intraneuronal transport. Left: Typical transport conditions in a control animal. NanoKTP (green cube) is spontaneously endocytosed and 24h after, ends up in an endolysosomal vesicle (light blue oval) of a PVN axon and is transported along a microtubule (MT, grey tube) in the anterograde direction (towards MT + end) by kinesins (green and yellow dimers to represent two different families, including kinesin 1) or in the retrograde direction (towards MT – end) by dyneins (blue dimers). Right: dynapyrazole impairment of microtubule-stimulated dynein activity. (B-I) Modifications of transport parameters in larvae bathed in dynapyrazole at 10 *μ*M concentration (condition DYNA), compared to control (bath without dynapyrazole, CTRL). For (B, D-I) bar plots display the parameter mean values ±s.e.m., while insets are distribution box plots of the same parameters. In (B) numbers in the bars are # of fields of view analyzed/# of animals analyzed, while for (C-I) they are # of trajectories/# of animals analyzed. (B) Fraction of moving particles, with directed motion. The treated larvae have 31% less moving particles than the control (*p* = 0.004). (C) Transport directionality. Pie charts of the percentage of purely anterograde (labeled A), purely retrograde (R), or bidirectional trajectories (B), for each condition. The proportion of purely A trajectories decreases in favor of purely R trajectories in treated larvae. Inset: fraction of retrograde transport in bidirectional trajectories (i.e. having at least one phase of R and one of A transport). A value of 1 means a purely retrograde trajectory, while a value of 0 signifies a purely anterograde one. This fraction of retrograde transport in B trajectories does not vary significantly (*p* = 0.738). (D-E) Retrograde and anterograde segmental velocities. Both R and A segmental velocities are not statistically different between the different conditions (*p* = 0.161 and 0.885, respectively). Trajectories with purely A and purely R movement were excluded from the analysis of R and A segmental velocities, respectively. (F) Pausing frequency. The pausing frequency of the treated larvae is 14% larger than that of the control (*p* = 0.004). (G) Pausing duration does not change under DYNA treatment (*p* = 0.261). (H) Retrograde run length is 37% shorter for the treated larvae (*p* = 0.004) compared to the control. Trajectories with purely anterograde movement were excluded from analysis. (I) Anterograde run length decreases by 28% in treated larvae, but there are no statistical differences between conditions (*p* = 0.084). Trajectories with purely retrograde movement were excluded from analysis.

**Figure 5:**
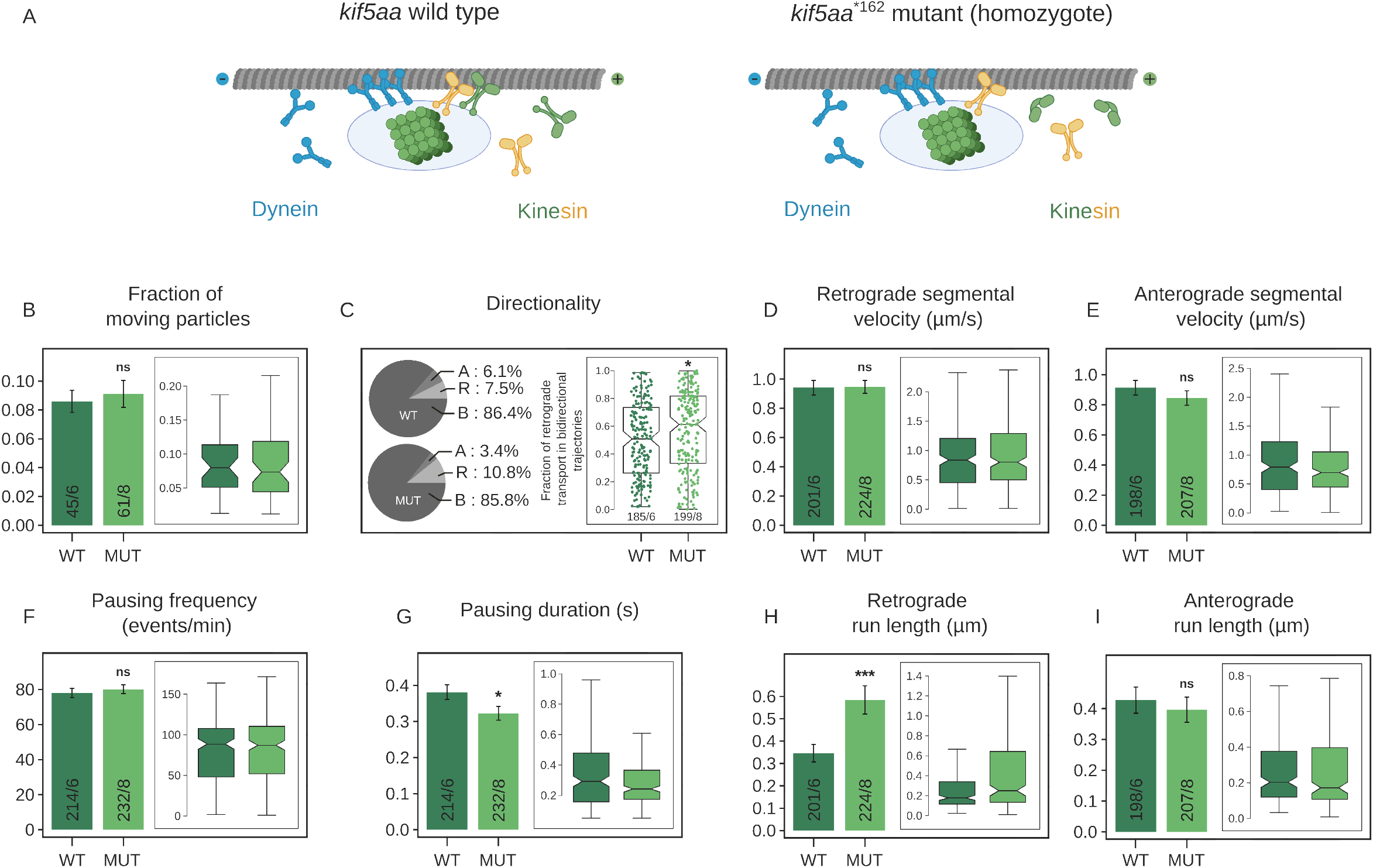
*kif5aa* mutation impairs intraneuronal transport in zebrafish brain. (A) Schematic representation of the mechanisms of intraneuronal transport. Left: Typical transport conditions in a wild type *kif5aa* sibling. The optically active nanoparticle (green cube) is spontaneously internalized in an endolysosomal vesicle (light blue oval) and is then transported along a microtubule (grey tube) in an anterograde motion by kinesins (green and yellow) or in a retrograde motion by dyneins (blue). Right: Impaired transport conditions by mutant kinesins (green shortened dimers). (B–I) Modifications of intraneuronal transport parameters in *kif5aa* larvae brain, depending on the larva genotype (WT: wild type or MUT: mutant homozygote). For (B, D-I) bar plots display the parameter mean values ±s.e.m., while insets are distribution box plots of the same parameter. In (B) numbers in the bars are # of fields of view analyzed/# of animals analyzed, while for (C-I) they are # of trajectories/# of animals analyzed. (B) Fraction of moving particles, with directed motion, showing no difference between WT and MUT (*p* = 0.83). (C) Directionality. Left: pie charts of the percentage of purely anterograde (A), purely retrograde (R), or bidirectional trajectories (B), for each condition. The proportion of purely A trajectories decreases in favor of purely R trajectories in MUT. Right: fraction of R transport in bidirectional trajectories. MUT shows a bias towards retrograde transport (*p* = 0.018). (D-E) Retrograde and anterograde segmental velocities do not vary between conditions (*p* = 0.896 and *p* = 0.181, for R and A respectively). Trajectories with purely anterograde or purely retrograde movement were excluded from analysis of retrograde or anterograde segmental velocities, respectively. (F) Pausing frequency, that does not vary (*p* = 0.574). (G) Pausing duration decreases by 16% (*p* = 0.046). (H) The average run length is 68% longer for MUT compared to WT (*p* = 0.00092). Trajectories with purely anterograde movement were excluded from analysis. (I) Anterograde run length does not vary (*p* = 0.303). Trajectories with purely retrograde movement were excluded from analysis.

### Key parameters for quantitative analysis of axonal transport with nanoKTP

Imaging data sets were further analyzed to extract quantitative parameters required to assess axonal transport defects. We first evaluated the precision of localization and applied a denoising strategy before applying the automated pipeline of transport analysis schematized on Fig. 1D. Trajectories of nanoKTP-labeled vesicles are inferred from consecutive positions of the nanoparticles (Materials and Methods) obtained as the position of the center of the best Gaussian fit to each SHG spot. Focusing on directed motions, we first filter-out “trajectories” corresponding to confined Brownian motions (possibly from particles in intercellular space) that are characterized by a bounded mean-square displacement.^39^

In the remaining trajectories, the detected positions are affected by a precision of localization noise whose 2D-standard deviation *σ_xy_* is expected to scale down with the number of detected photons *N*, as 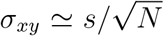, in the case of a photon shot noise-limited process, where *s* is the standard deviation of the microscope objective experimental point spread function (PSF_exp_). We estimated *N* after having determined a conversion factor between the 8-bits grey level of the hybrid photomultiplier detector (Supporting Data S3 and Figure S3), and we inferred s ≈ 355 nm from the smallest SHG spots (Supporting Data S4 and Figure S4A).

We measured *σ_xy_*(*N*) from videos of static nanoKTP (dropcasted on a glass coverslip) in the same acquisition conditions as for the intraneuronal transport (20 fps, in the cage incubator). Supporting Figure S4A displays one SHG spot from Supporting Video S3 and Gaussian fits of two orthogonal cross-sections of this spot. Figure S4B is a scatter plot of Gaussian fit centers positions of the same nanoKTP spot in consecutive frames. The precision of localization of this spot is given by the 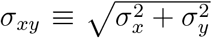, where *σ_x_* and *σ_y_* are the standard deviations in *x* and *y* directions.

We then considered an ensemble of SHG spots with different *N* and plotted on Supporting Figure S4C *σ_xy_*(*N*). The smallest experimental precision is 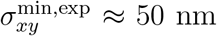, obtained for *N* = 440 photons, while the theoretical minimal value for the same *N* is 10 nm (Supporting Data S4). We attribute this ≈ 40 nm excess of noise mostly to mechanical vibrations. At the other extremity, the worst localization precision of 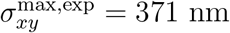 correspond to the dimmest spot. Altogether the average value is 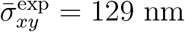.

Considering the blinking of the SHG signal due to the vesicle rotational mobility, a spot that appears bright in one frame and can be localized with 50 nm precision, may also become very dim a few frames after and be localized with a precision of only ≈ 370 nm. Such variations along one trajectory may induce artifacts in transport parameters quantification. For instance, in the extreme case of two consecutive positions separated by 370 nm, we cannot discriminate between a static faint particle and a very fast displacement, at an instantaneous velocity of 0.370/0.05 ≈ 7.4 *μ*m/s, which is very unlikely as the average reported velocity in zebrafish neurons is in the 0.4-1.2 *μ*m/s range.^40^

In order to mitigate the impact on transport parameters values of a low SHG intensity (associated to a large localization uncertainty), we regularized the trajectories by a nonlinear total variation-based noise removal algorithm^41^ (Fig. 1D). As the localization uncertainty introduces variations of the instantaneous velocity larger than the one expected from the true vesicular motion (constituted of phases of almost constant velocity, sparsely interrupted by much slower motion or complete pauses) we implemented a regularization algorithm (see Materials and Methods) that infer the positions’ ground truth by minimizing the acceleration (velocity changes), under a constraint related to the average experimental precision of localization of 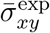 previously determined separately.

Based on the straight pattern of PVN axonal projections, endolysosomal trajectories are expected not to deviate strongly from straight lines. Hence, at this stage of the analysis, we excluded the trajectories that were not properly fitted by a third order polynomial law (Materials and Methods), which represented 17% of them.

The following step of the automated analysis consists in parsing the regularized trajectories in phases of “slow” (further called Stop) and “fast” (Go) motions (Fig. 1D). To this aim we apply the same heuristic method as in our previous work. ^22^ Briefly, we calculate at each trajectory position i a confinement ratio *R_i_* on a sliding window of 6 consecutive points width, defined as the net distance between the first and last point divided by the sum of the length of the two segments. *R_i_* = 1 corresponds to straight line, and therefore a Go phase, while *R_i_* = 0 is associated to a zero net motion, hence a Stop phase. To attribute a phase state to all other intermediate values of *R_i_*, we searched for a confinement ratio threshold *R*_th_ emerging from the datasets. To this aim we examined the distribution of *R_i_* for all trajectories of all conditions (including heterozygote larvae) in *kif5aa* larvae (Supporting Data S5 and Figure S5) and noticed that it has a minimum at 0.64. We then decided to consider this minimum as the threshold to parse the phase states: *i*^th^ point with *R_i_* > *R*_th_ = 0.64 is in a Go phase, while for *R_i_* ≤ *R*_th_ the vesicle is in a Stop phase.

For the quantitative analysis of the transport, we calculate two general parameters and six metrics associated to individual trajectories, distinguishing retrograde from anterograde displacements according to previously described criteria (*i.e*. the sign of the displacement along (O*x*) and the eye side). The two general parameters are the fraction of moving particles detected in each video (*i.e*. in each field of view), and the transport directionality evaluated as the proportions of purely retrograde (R), anterograde (A) or bidirectional trajectories (B). In the case of bidirectional trajectories, we refine the analysis by calculating the fraction of retrograde Go phase over all types of phases (calculated from the traveled distances).

We then characterize each trajectory by the six metrics inferred from the parsing: retrograde and anterograde segmental velocities (SV, average velocities during R and A Go phases only), the pausing frequency (PF, number of Stop phases per minutes), the average pausing duration (PD), the retrograde and anterograde run lengths (RL, average distances traveled during R and A phases).

In order to check that the mere injection of nanoKTP does not impact axonal transport, we investigated lysosomal transport, in wild type and injected conditions. This was assessed by exposing the larvae with LysoTracker Red, a live labeling dye of lysosomes. To quantify the axonal transport of LysoTracker-positive vesicles, we used exactly the same pipeline of recording and analysis than for nanoKTP, here taking advantage of the two-photon absorption of the dye. Despite a lower signal-to-background ratio for LysoTracker than for nanoKTP and bleaching, we were able to extract enough reliable trajectories to compare the transport parameters with or without nanoKTP, and did not find any difference between these two conditions as shown in Supplementary Figue S6. Hence our method, with its sparse labeling of tracked compartments (as shown on Fig. 2E-F), does not only have no measurable impact on lysosomal transport parameters, but it also provides a larger number of reliable trajectories, owing to the large signal-to-background and perfect photostability of the SHG signal.

This LysoTracker Red labeling experiment gave us also the opportunity to evaluate the colocalization of nanoKTP with late endosomes or lysosomes, which are the endolysosomal compartments positive to LysoTracker. We only considered live colocalizations of nanoKTP having directed motion and found that about 56% of them colocalize with LysoTracker Red, as shown in Supplementary Figue S7. This observation is consistent with the timeline of the endocytosis cycle, since we record nanoKTP motion 24h after their injection, at a stage where the majority of the up-taken nanoparticles is expected to have reached late endosomes and lysosomes. The remaining fraction of 44% may correspond to nanoKTP that are in less acidic endosomal compartments, with lower LysoTracker Red signal. Hereafter, we name the mobile nanoKTP containing compartments “endolysosomal vesicles”, shortened sometimes in “vesicles”.

In the following we show that the general parameters and trajectory metrics defined above and evaluated for nanoKTP-labeled endolysosomal vesicles, are sensitive enough to measure the impact of drugs inhibiting the motors, or loss-of-function mutations, starting with dynein (using either a mutant of dynein heavy chain 1 or a dynein inhibitor) and continuing with the kinesin 1 mutant *kif5aa*.

### Mutation or inhibition of dynein motors affects endolysosomal transport *in vivo*

We first consider the *mw20* zebrafish transgenic line with a point mutation of *dync1h1* (*dync1h1*^mw20^) coding for heavy chain 1 of the dynein motor complex. This mutation creates a premature stop codon leading to a non-functional dynein devoid of the tail (that binds to the microtubule, see Figure 3A) and of the domains associated to ATPase activity. ^31^ Despite this mutation, the maternal contribution of wild-type *dync1h1* mRNA and protein leads to larvae survival at least during 8 days, which fits our experimental plan.

Fig. 3(B-I) display axonal transport properties in homozygous (HOM) mutant, heterozygous (HET) or wild-type (WT) siblings. We use the fraction of moving particle in each field of view (of similar morphology) as a first parameter to quantify the overall impact of the mutation on the transport. In homozygous *dync1h1*^mw20^ mutant data set we detected only one particle having a directed motion corresponding to an estimated mobile fraction of 0.01%, in contrast to the wild-type and heterozygous larvae in which this fraction respectively reached 4.8% and 5.2% (Fig. 3B). Furthermore, we observed no differences between WT and HET larvae in any of the other transport parameters (Fig. 3(C-I)). We attribute these results to the fact that most dynein motor complexes are inactive even in normal conditions as they need to be associated to activators (as shown in Redwine *et al*.^42^), making it possible to maintain a normal transport with only half of the WT concentration of dynein heavy chain 1, as long as this half forms the same concentration of active complexes than in WT conditions. Such a phenomenon may explain the absence of transport parameter modifications in HET *dync1h1*^mw20^ larvae compared to WT siblings. Regarding the vesicle directionality (pie plots of Fig. 3C), the dominant category of bidirectional motion within single trajectories is consistent with observations reported in experiments using high-resolution analysis like ours,^23^ and most likely reveals a tug-of-war situation of motors pulling in opposite direction alternatively, leading however to a net displacement which is either anterograde or retrograde. As a refinement, we also investigated the fraction of retrograde transport within the population of bidirectional trajectories only (inset of Figure 3C). This fraction would be equal to 1 for a trajectory made of purely retrograde segments. It did not show differences between WT and HET.

Dynein being the only retrograde motor, a strongly modified retrograde transport was expected in *dync1h1*^mw20^ mutants, but the anterograde transport was also disrupted. Indeed, this observation is consistent with several reports in which the interruption of retrograde transport, in particular *via* a mutation on dynactin (complexing with dynein to activate it), impacts the anterograde transport too.^43–45^

As the endolysosomal transport phenotype of *dync1h1*^mw20^ homozygous larvae is severe, we decided to challenge our axonal transport measurement method with another model of retrograde transport impairment. Zebrafish larvae were then treated with a dynein inhibitor for which concentration can be adjusted to produce moderate effects. To this aim, we selected dynapyrazole-A (DYNA), a new conformational isostere of the first-in-class dynein inhibitor ciliobrevin, that has a higher selectivity in blocking only dynein’s microtubule-stimulated activity (Figure 4A) and not the ATPase one. ^33^

In *in vitro* assays DYNA strongly inhibits dynein’s microtubule-stimulated activity at micromolar concentrations. In absence of data available *in vivo*, we first immersed Zf larvae in different concentration (10 and 15 *μ*M) of dynapyrazole for 25 to 45 min at 28°C in a bath. The larvae were then washed and observed to monitor the appearance of potential toxicity effects right after or 1 h post exposure. Thirty minutes incubation at 15 *μ*M lead to severe malformations (body curvature) detected 1 h post exposure, which evolved into larvae death (data not shown). As there was no toxicity at 10 *μ*M (25 min exposure), the following experiments were performed at this concentration. Figure 4B-I display the effect of DYNA on the axonal transport properties. We use the fraction of moving particle in each field of view (of similar morphology) as a first parameter to quantify the overall impact of DYNA on the transport. In untreated control sample, this fraction is about 6.7%. while it decreases to 4.6% upon DYNA exposure, representing a 32% decrease (Figure 4B). As DYNA is not known to impact endocytosis, we attribute this reduction to the inhibition of dynein’s microtubule-stimulated activity. Our observations are in agreement with the transport impairment of fluorescently labeled lysosomes initially reported *in vitro*.^33^ We then questioned whether this dynein inhibition modifies the preferred direction of transport. Pie plots of Figure 4C displays the proportions of purely retrograde, anterograde or bidirectional trajectories for both control and DYNA conditions. The total fraction of purely directional trajectories is slightly reduced by DYNA (16.3% in control to 13.5% in DYNA). In a counter-intuitive way, within this fraction, the one of purely anterograde transport decreases by 42% from 8.4% (control) to 4.9% (DYNA), while purely retrograde transport stays almost constant evolving from 7.9% (control) to 8.6% (DYNA). The fraction of retrograde transport within the population of bidirectional trajectories only (inset of Figure 4C) has a very broad distribution, indicating that even bidirectional trajectories are polarized (with a slight bias towards R direction as evidenced by the value of the median of 0.57), leading to a net displacement in either R or A direction. We observed that the fraction of retrograde transport in these bidirectional trajectories does not vary significantly between control and DYNA, but we noticed however that its distribution slightly narrows from control to DYNA, which is the sign of a slight decrease of polarity. This conclusion is further confirmed by the significant increase of 19% of the number of direction reversals per minute within a trajectory (Supporting Figure S8A).

We then focused our attention on specific parameters of the retrograde transport. We did not detect significant variations of the R and A segmental velocities (Figure 4D-E) which values are around 0.9 *μ*m/s for both directions, consistent with values reported for axonal transport of various endosomes in Zf larvae motor neurons.^12,40^ On the contrary we observed a significant increase of about 15% of the pausing frequency (Figure 4F) with DYNA (from 70.4 to 80.7 events/min), accompanied with a trend of augmentation of the pausing duration, all types of pauses included (Figure 4G, from 0.30 to 0.32 s). We investigated whether this trend could be due to a sub-population of pauses, considering the ones between either anterograde, opposite directions, of retrograde phase of motions (Supplementary Figure S9A-C). None of these categories revealed any differences of pausing duration for DYNA treated Zf compared to control.

Regarding the spatial parameters, we measured a significant decrease of the retrograde run length by 37% from 622 to 390 nm (Figure 4H) concomitant with a trend of decrease by 28% of anterograde run length from 570 to 409 nm (Figure 4I). Our observations are in agreement with previous reports in which the reduction of cytoplasmic dynein concentration was observed to impair the transport parameters^46^ with a similar reduction of anterograde RL, indicating that dynein is also involved in the activation of anterograde transport.

We then considered the perturbation of motors driving the anterograde transport in PVN neurons of Zf brain. To this aim we investigated the effect of a change of concentration of the dominant kinesin motor in the brain, kinesin 1, as induced in a mutant zebrafish.

### Precise measurement of slight endolysosomal transport defects induced by a kinesin 1 motor disruption

Anterograde vesicle displacement along microtubule is mediated by kinesin superfamily proteins^4^ that also play a key role in the organism development.^47^ To validate our model of *in vivo* detection of fine axonal transport impairment caused by abnormal kinesin concentration, we monitored nanoKTP axonal transport in the *kif5aa* zebrafish transgenic line. This mutant generated by Auer *et al*., ^32^ was engineered to introduce a premature stop in the motor part of Kif5aa molecule, hence preventing its complete synthesis and therefore its function, leading to abnormal phenotype that includes blindness and failure to inflate the swim bladder, eventually resulting in death around 10 dpf.

Moreover, this mutant is relevant to our experimental demonstration as *kif5aa* expression was shown to be specific to neurons,^48^ and in wild-type siblings of *kif5aa* transgenic line at 3 dpf, *kif5aa* is expressed in the whole optic tectum, including PVN neurons (refer to Fig.1B of ref.^32^). *Kif5aa* mRNA is also down-regulated in mutant larvae probably due to non sense-mediated decay of the mutant transcript containing a premature stop codon that leads to the translation of a non functional truncated protein. Considering also that *kif5aa* is not maternally expressed in the germline,^48^ and therefore Kif5aa not maternally contributed, we expect that, in *kif5aa* mutant, the level of functional Kif5aa in axons of PVN (the brain region we studied) is smaller than in wild-type siblings. However, other types of kinesins could compensate the decrease of functional Kif5aa and normalize the transport, including the protein products of the other four *kif5* paralogs present in Zf encoding for kinesin 1 heavy chain isoforms. ^48^ This is in contrast to the previous situation where dynein represents the only motor driving retrograde transport.

In *kif5aa* larvae (Figure 5A) we evaluated the same axonal transport parameters as in dynein inhibition experiment, comparing mutant (MUT) to wild-type siblings (WT). We did not detect differences in the fraction of moving particles between these two conditions in axons of the PVN (Figure 5B). These observations differ from previously published results conducted in the axons of retinal ganglion cells of *kif5aa* mutants showing an increase of the mobile fraction of small vesicles. ^32^

Then, we investigated the transport directionality (pie plots of Figure 5C). In mutant larvae, while the total fraction of purely directional motion only slightly increases from 13.6% to 14.2%, the fraction of pure A trajectories is only 3.4% in the MUT to the benefit of pure R trajectories which increases from 7.5% to 10.8%. As for the study of DYNA effect, we also focused on the remaining directionality in the largest fraction of bidirectional trajectories. We evaluated the fraction of retrograde transport in this sub-population and found that it is broadly distributed (inset of Figure 5C) indicating that bidirectional trajectories are largely polarized. Moreover, this fraction also displays a median shift from 0.51 to 0.61 between WT and MUT, which means that bidirectional trajectories are biased towards the retrograde direction in mutants.

Furthermore, as observed in the case of dynein inhibition with DYNA, in *kif5aa* mutant we did not detect significant changes of both A and R segmental velocities that remain of 0.8-0.9 *μ*m/s (Figure 5D-E). The pausing frequency does not vary statistically neither between WT and MUT, but we observe a decrease of the pausing duration (all types of pauses gathered) of about 15% from 0.382 to 0.323 s. Decreases were also observed for all pause subcategories (i.e. between two identical directions or two opposite directions of motion), but none of them was individually significant (Fig.S9D-F). Moreover, we observe a significant increase of the retrograde RL by 68% from 347 to 585 nm between WT and MUT, without effect on the anterograde RL (Figure 5H-I).

Altogether, the directionality study and endolysosomal transport parameters in *kif5aa* larvae indicate that reducing Kif5aa concentration biases the direction of transport towards the retrograde direction and favor longer R run lengths. These results are compatible with the tug-of-war model of intraneuronal transport,^23^ stating that (i) both types of R and A motors (dynein and kinesins respectively) are present on the cargo, (ii) that they are both attached to the microtubule pulling in opposite directions, and (iii) that the direction of actual motion is dictated by the set of active bound motors pulling the strongest, which is likely to be the dynein motors in our situation of reduced amount of Kif5aa kinesin 1 motor.

The absence of modification of anterograde velocity (Figure 5E) and more surprisingly of anterograde run length (Figure 5I) in *kif5aa* mutants is in favor of a compensation mechanism. In Zf larvae such compensation can occur by other Kif5 motors to ensure anterograde motion, or by another kinesin motor complex of the superfamily, such as kinesin 2 motor complex including Kif3a/3b/Kap3 and Kif3a/3c/Kap3, ^49^ which is known to drive vesicle axonal transport too.^23,50^ In the case of inhibitor experiments, as dynein heavy chain 1 and 2 isoforms are the only motors ensuring retrograde transport, and as they were reported to be both inhibited by dynapyrazole A *in vitro*,^33^ no motor substitution is expected to compensate their inactivation. However, at the dynapyrazole concentration used, the retrograde transport is not fully halted, and the overall transport displays only a reduced retrograde run length (Figure 4H) and increase in pausing frequency (Figure 4F), with no modification of velocity (Figure 4E). These observations may reveal a limited intravital penetration of dynapyrazole A, but we could not test this hypothesis because of the toxicity observed for higher concentration (possibly due to dynein 2 concomitant inhibition).

### NanoKTP polar angle fluctuations measurement

Finally, we took advantage of the blinking of the SHG signal from nanoKTP to reveal modifications in the endosome/lysosome-motor-microtubule binding in *kif5aa* mutants. Indeed, this blinking reflects changes of orientation of the vesicle relative to the microscope objective axis (Figure 1C), as the nanocrystal is presumed immobile relative to its embedding vesicle. ^29^ To quantify this fluctuation, we first reverted from 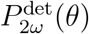 (the detected SHG power at frequency 2*ω*), the polar angle *θ* between the direction of light propagation and nanoKTP *c*-axis (which is also the direction of the equivalent SHG-emitting dipole, see Figure 1C). To this aim we calculated the efficiency *η*(*θ*) of the SHG power signal collection through the microscope objective in function of *θ*, and obtained the general form *η*(*θ*) = 3/8*π*(*B* sin^2^ *θ* + *C*), where *B* and *C* are parameters depending only on the microscope objective NA (Supporting Information Data S10). SHG detected power differs from SHG collected one *P*_2*ω*_ by a constant coefficient taking into account optical losses and the detector quantum efficiency. Figure 2A shows that all SHG signals from nanoKTP-labeled vesicles with directed motion vary between almost zero power 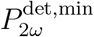 and a maximum 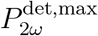 that depends on the particle size. We assumed that these extreme power values correspond to *θ* = 0° and 90° respectively, and then inferred *θ* at each trajectory point on from mapping the detected power to *η*(*θ*) by 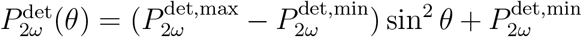. Note that we are not able to extract the azimuthal angle defining the orientation of the SHG dipole in the sample plane, as we are not resolving the polarization of the collected light.

Then in order to quantify the fluctuations of vesicle orientation, we first applied a Savitzky-Golay filter to SHG intensity time traces in order to mitigate the high sensitivity of the intensity-to-polar angle inversion process to a large intensity fluctuation close to the extrema. We determined *θ*(*t*) from the filtered intensity time trace and calculated its standard deviation *σ_θ_* in each phase of motion within a trajectory, from which we inferred an average value 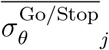 per trajectory *j*. We focused our analysis on *kif5aa* data set. Figure 6A displays the distribution of these values for all the trajectories of WT and MUT conditions. We notice that in the MUT case 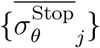 distribution (simply noted 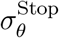 after) is significantly larger by 8% than the one of 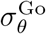 and that there is no difference in the WT case. Comparing now 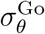 and 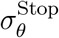 between WT and MUT, we did not observe differences of orientational fluctuations between MUT and WT (Figure 6B). To our knowledge, this is the first time such fluctuations of endolysosomal vesicle orientation are reported *in vivo*. In the case of the mutant, our data are in agreement with observations performed *in vitro* in cultured PC12 rat neuronal cell line showing larger orientational fluctuation of the vesicle in Stop than Go phases, as detected from differential interference contrast signal from embedded gold nanorods. ^28^ However, in primary cultures of dorsal root ganglion rat neurons, Kaplan et al.,^29^ did not observe any differences of the rotational lability (another measure of the rotational fluctuation) between the Stop phases and Go phases, like in the case of *kif5a* WT siblings.

**Figure 6:**
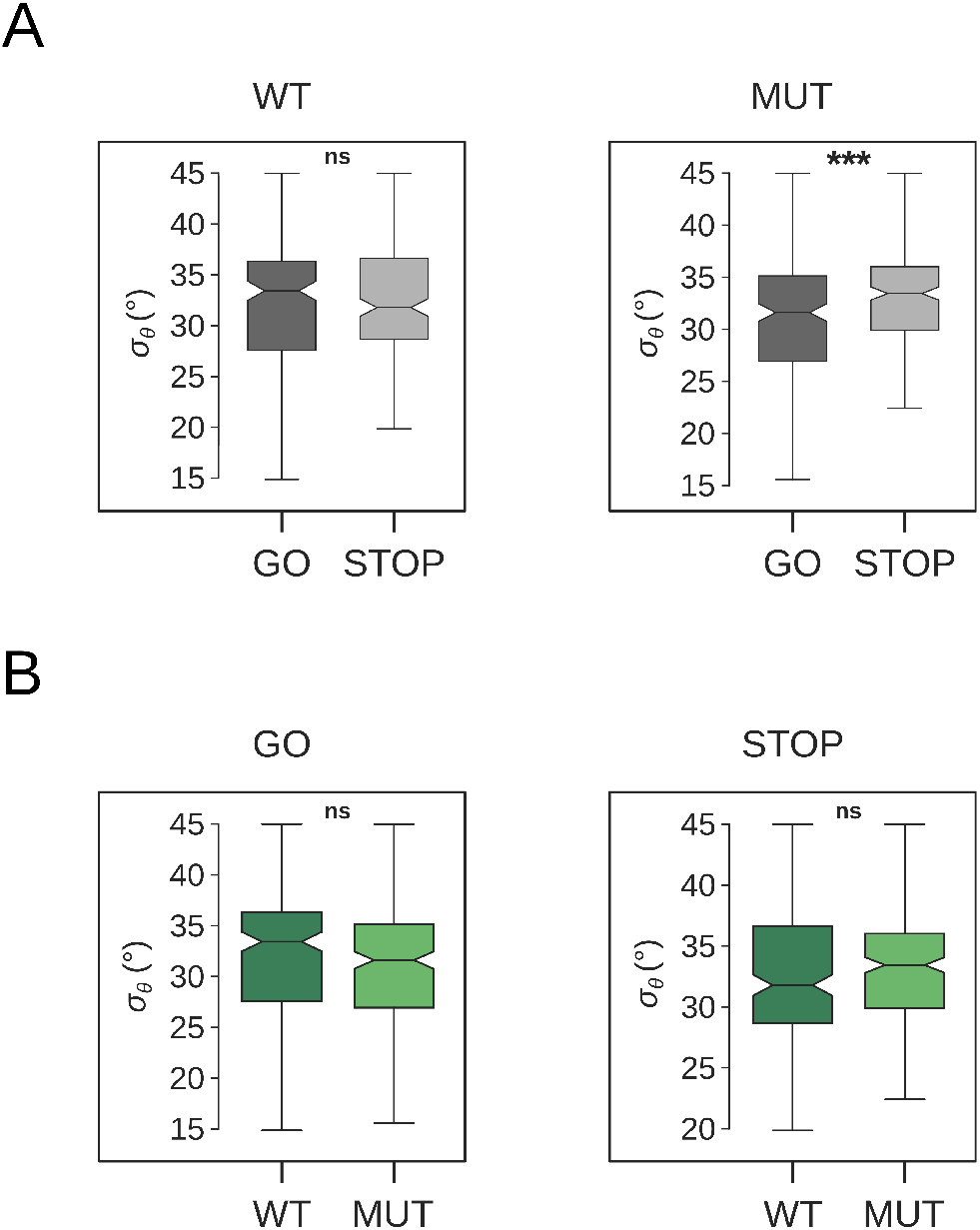
Fluctuation of the orientation of nanoKTP-labeled endosome vesicle during Go and Stop phases of motion in neurons of the brain of *kif5aa* Zf larvae, as quantified with the standard deviation *σ_θ_* of the polar angle *θ*. (A) Comparison of the fluctuations between the Go and Stop phase for the two conditions, showing that *σ_θ_* has no difference between Go and Stop phases (*p* = 0.99), but the fluctuations are larger in the Stop (average of 33.1°) compared to the Go (30.6°) in the mutant (*p* = 1.5 × 10^-4^). (B) Comparison of *σ_θ_* between the two conditions in the different phases, showing no difference between WT and MUT in the Go (*p* = 0.06) and Stop (*p* = 0.07) phases.

In order to check the consistency of our observations in different conditions, we also computed the vesicle orientation fluctuations for the dynapyrazole dataset (Supporting Figure S12). We observed similar distributions and median values of *σ_θ_* than in *kif5aa* data, with larger 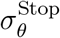 than 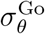 for both control and DYNA. Here, the inhibition of functional dyneins by dynapyrazole leads to an increase of 14% of *σ_θ_* in the Go phase, and no changes in the Stop ones (Figure S12B).

## Conclusions

In this work we described a novel method for *in vivo* quantification of axonal transport in neurons of the CNS, based on the measurement of a large set of parameters that are able to reveal transport impairments finely. This method relies on intravital fast two-photon microscopy recording of the motion of axonal endolysosomal vesicles, thanks to their labeling with nonlinear nanocrystals of KTP, as a first step, followed by a high-throughput data processing extraction of endosome trajectories and associated transport metrics. We established this approach *in vivo* by using the zebrafish model organism, exploiting its translucency and genome editing possibilities. We achieved sparse labeling of endolysosomal vesicles traced in the axonal compartments of CNS neurons through microinjection of the nanoKTP in the optic tectum of Zf larvae. About 90% of the injected larvae had, in the region of interest, a concentration of nanoKTP well suited to isolate single trajectories after the nanocrystals spontaneous axonal uptake.

This easy to apply original approach contrasts with the use of transient knock-in consisting in the injection of nucleic acid sequences encoding fluorescent proteins and sorting of the larvae presenting the relevant phenotype. Moreover, the nanoKTP are well tolerated by the Zf larvae and exhibit a large SHG signal that is perfectly photostable, in contrast to conventional molecule-based fluorophores that bleach during the recording. Our approach allowed us to reach an average resolution of 129 nm at 20 frames per second, which is a photon budget-limited compromise between high spatial resolution and fast acquisition. This spatiotemporal resolution achieved in our method has no equivalent in *in vivo* studies of axonal transport measurements to date.^6^ Therefore it constitutes a major advance to precisely quantify slow/Stop phases-related parameters and to detect altered cargo pausing associated to abnormal neuronal activity. In addition, the intracerebral injection in the optic tectum of the Zf brain led to an efficient uptake of nanoKTP localized in the axons of the periventricular neurons, allowing to study independently kinesin or dynein driven directions of motion.

To demonstrate the sensitivity of our *in vivo* assay, we probed the impact of reducing the concentration of the active molecular motors on the transport dynamics. To this aim we reduced the concentration of dynein (using dynapyrazole, the best-in-class dynein inhibitor) or kinesin 1 Kif5aa (using *kif5aa* loss-of-function zebrafish mutant line) which is a member of kinesin 5Aa family, abundantly expressed in neurons. We observed differential overall effects of these modifications: dynapyrazole reduces by 32% the mobile fraction of nanoKTP directed motion, proving its efficiency as a dynein inhibitor in Zf larvae brain. No difference in this fraction was detected between wild type and mutant *kif5aa* larvae. Dynapyrazole also consistently reduces the retrograde run length by 37%. *Kif5aa* mutants showed a 68% increase of the retrograde run length, instead of a reduction of the anterograde run length. To interpret this observation, we hypothesize that other kinesins might replace Kif5aa and maintain the anterograde transport. As such, kinesin 2 is one possible candidate as it binds more loosely to the microtubule,^51^ which could explain the larger orientational fluctuations of the endolysosomal vesicle-containing nanoKTP in mutant *kif5aa*. However, further experiments are required to validate this interpretation.

In our current assay, the spontaneous endocytosis of bare nanoKTP in axonal compartments could be enhanced by conjugating transferrin or rabies virus glycoprotein peptides to nanoKTP. Future developments will also extend the use of our method to other neuronal cell subtypes and/or later stages of development. Taking advantage of the low absorption of tissues in near infrared excitation wavelength, and the possibility to easily tune excitation wavelength minimize the tissue auto-fluorescence contribution, our method open prospects to image more complex and thicker biological model, such as juveniles or adults zebrafish mutants devoid of pigmented cells. ^52^ These approaches will provide a way to study axonal transport in physiological conditions at different stages of development in normal and pathological situations. The latter include influence of intrinsic (*i.e*. mutations inducing transport defects) or extrinsic (*i.e*. temperature, water quality, chemical exposure…) factors.

Finally, our assay could also be valuable to develop biomedical models reproducing human nervous system alterations quantitatively assess the link between axonal transport impairment and disease outcome. In particular, it would be interesting to reproduce our experiments in Zf larvae infected by neurotropic viruses,^53–55^ in transgenic lines used as models of neurodegenerative diseases,^56–58^ or upon axonal photoablation^59^ to precisely measure the axonal transport impairments in these situations. In case of detection of a significant impairment of axonal transport, our assay could be harnessed to screen therapeutic compounds able to rescue the defect.

## Materials and Methods

### Zebrafish

Zebrafishes (*Danio rerio*) were maintained at 28°C on a 14 h light/10 h dark cycle. The mutant line *kif5aa*^*162^ was generated as previous described.^32^ *dync1h1*^mw20^ mutant line was described by Insinna *et al*. ^31^

#### Ethics statement

Zebrafishes were housed in the animal facility of our laboratories which were built according to the respective local animal welfare standards. All animals were handled in strict accordance with good animal practice as defined by the European Union guidelines for the handling of laboratory animals (http://ec.europa.eu/environment/chemicals/lab_animals/home_en.htm). All animal procedures were performed in accordance with French and European Union animal welfare guidelines with protocols approved by committees on ethics of animal experimentation of Sorbonne Université (APAFIS#21323-2019062416186982) and Université Paris-Saclay. All animal work was also approved by the Direction of the Veterinary Services of Versailles, France (authorization number C78-720), and experimental protocols were approved by the INRAE institutional ethical committee “Comethea” (APAFIS#22160-2019092615256610).

### Genotyping of *dync1h1*^mw20^ and of the *kif5aa*^*162^ mutant alleles

For *kif5aa*^*162^ genotyping the following primers were used (5’ to 3’): kif5aa geno fwd: GTTCACAGATTGTGATGTCTGTG, kif5aa geno rev: TGGAGGATG-GAGAAATGAT-GACA. After PCR amplification from genomic DNA the 400 bp long amplicon was digested with NcoI. The wild-type allele is digested into two fragments of 240 bp and 160 bp length, respectively. The mutant homozygote alleles is not digested and show a single band at 387 bp or 390 bp, while the heterozygote genotype is characterized by three bands at 160, 240 and 390 bp. For *dync1h1*^mw20^ genotyping the following primers were used (5’ to 3’): mw20 geno fwd: CACGAGGAGCTCTACAAGTGG, mw20 geno rev: GAACAGGTTG-GCGTAGTGGT. After PCR amplification from genomic DNA the 750 bp long amplicon was digested with RsaI. The wild-type allele is digested into two fragments of 350 bp. The mutant homozygote alleles is not digested and show a single band at 750 bp, while the heterozygote genotype is characterized by two bands at 750 and 350 bp.

#### KTiOPO_4_ nanocrystal synthesis and size separation

KTP nanoparticles were synthesized following a protocol adapted from Ref. ^26^ In a typical experiment, a titanium alkoxide solution acidified with HCl is first mixed with a solution of monobasic potassium phosphate leading to the formation of a precipitate. The solution is then neutralized up to pH=6 through the addition of K_2_CO_3_. The precipitate is then recovered by centrifugation, washed several times with water and dried. The powder is then thermally treated to ensure KTP phase formation with an optimal temperature of 700° for 2 hours. The powder is then washed three times by centrifugation with deionized water and the pellet redispersed in water leading to the colloidal suspension of KTP nanocrystals.

The hydrodynamic particle size for this pristine suspension was estimated to be 230 nm by dynamic light scattering (DLS, volumic fraction). As previous characterization have shown a significant size dispersion, ^26^ we decided to apply a size selection procedure to narrow the size distribution. To this aim, the pristine suspension was first centrifugated at an acceleration of 8,000 g for 5 min. Then, the supernatant was extracted and centrifuged at 60,000 *g* for 5 min. The supernatant was discarded and the pellet was redispersed in pure water leading to the colloidal suspension used in this work. DLS analysis of this solution gave a cumulant *Z*-average particle size of 126 nm (volumic fraction analysis), confirmed by nanoparticle tracking analysis (Supporting Data S1) yielding a value of 123.8±2.4 nm.

#### Zebrafish husbandry and microinjection of nanoparticles

Wild-type (AB strain) were raised in the IERP fish facilty at INRAE Jouy-en-Josas, France. After spawning, eggs were collected and incubated at 28°C in fish system water supplemented with 0.3 *μ*g/ml methylene blue. Phenylthiourea (PTU) was added at 24 hours post-fertilization to inhibit synthesis of melanin. Before injection, larvae were anaesthetized with eugenol diluted at 0.0075% in fish system water supplemented with methylene blue and PTU as described above. Larvae were injected in the left brain hemisphere with 5 nl of nKTP solution using pulled borosilicate glass microcapillary (GC100F-15, Harvard Apparatus) pipettes under a stereomicroscope (MZ10F, Leica Microsystems) with a mechanical micromanipulator (M-152; Narishige), and a Femtojet microinjector (Eppendorf).

To investigate the transport properties of lysosomes and the colocalization of moving nanoKTP with these compartments, 4 dpf zebrafish larvae were exposed during 1 hour to LysoTracker Red DND-99 (Ref.L7528, Invitrogen/ThermoFisher) diluted at 5 μM in fish system water supplemented with methylene blue and PTU. Larvae were rinsed briefly before mounting for live imaging.

#### Immunostaining

Zebrafish larvae were fixed overnight at 4°C in PBS 0.01 M + 4% formaldehyde. Larvae were rinsed three times for 5 minutes with PBS + 0.1% tween. All the following steps were performed at room temperature (RT) unless indicated otherwise. Larvae were dehydrated in ascending methanol series (20%, 40%, 60%, 80% in H20) during 15 minutes for each step. Larvae were incubated in 100% methanol overnight at 4°C. Dehydrated larvae were rehydrated in descending methanol series (80%, 60%, 40%, 20% in water) during 15 minutes for each step. Samples were permeabilized by immersion in PBS + 0.5 *μ*M CaCl_2_ + 1 mg/ml collagenase (C9891, Sigma-Aldrich) during 15 minutes. Larvae were rinsed three times during 5 minutes in PBS + 0.2% triton X100. Larvae were then blocked in PBS + 5% triton X100 + 10% DMSO + 10% horse serum + 0.05% sodium azide for 5 hours at 37°C. Mouse anti-acetylated tubulin (T7451, Sigma-Aldrich) and rabbit anti-GFAP (ZO334, Dako) were both diluted at 1/500 in PBS + 5% Triton + 10% DMSO + 10% horse serum + 0.05% sodium azide and applied 2 days at 37°C. After repeated washes in PBS + 5% triton + 10% DMSO + 0.05% sodium azide, samples were incubated overnight at 37°C with Alexa Fluor 488 goat anti-mouse (A11001, ThermoFisher) and Alexa Fluor 594 goat anti-rabbit (A11012, ThermoFisher,) diluted at 1:500 in the same solution than first antibody was applied. Larvae were rinsed several times with PBS + 0.1% tween. Before imaging, larvae were cleared by incubation in RIMS overnight. ^60^ Larvae were mounted under #1 coverslips in RIMS supplemented with 0.8% low-gelling agarose.

#### Non-linear microscopy live imaging and immunostaining imaging

Non-linear live imaging of zebrafish larvae was performed with an upright Leica SP8 two-photon microscope using a HCX IRAPO 25X/0.95NA water immersion objective (Leica Microsystems). Anaesthetized larvae were transferred to 3% agarose casts containing wells allowing to position larvae. Larvae were then embedded in 0.5% low-gelling agarose. Orientation of larvae was adjusted to dorsal view before complete gelation. Fish system water + methylene blue + PTU + eugenol was added on embedded larvae. NanoKTP were excited at 1040 nm with a Chameleon Vision II laser (Coherent). A quater-waveplate was used to convert the linear excitation polarization into a circular one. The emitted photons were detected in a non-descanned (NDD) pathway with a Gallium Arsenide Phosphorous-based hybrid detector (HyD, Leica), having 50% quantum efficiency at 520 nm wavelength, and a maximum counting rate of 60 Mcounts/s in the conventional photon counting mode, extended to 300 Mcounts/s in the “standard” mode that we used, and that compensate for the nonlinearity of conventional counting. We put a 525/50 nm bandpass (green channel) in front of the HyD. Temperature was maintained constant during and between experiments by using an incubator box combined with a precision air heater (The Cube, Life Imaging Service). All experiments were performed at 24°. For the live imaging of LysoTracker Red with two-photon excitation, we tuned the pulsed laser wavelength at 1030 nm, as 1040 nm corresponds to a minimum of the two-photon absorption of this dye. ^61^ This had no impact on nanoKTP SHG detection as its spectrum, now centered on 515 nm with 10 nm FWHM, still falls within the 525/50 nm bandpass filter of the green channel. We detected the LysoTracker Red emission (red channel) on a similar HyD as the green, placed on the transmission path of the beamsplitter (edge at 560 nm wavelength) separating green and red channel, and preceded by a 585/40 nm bandpass filter.

Immunostained samples were acquired by confocal and 2-photon microscopy using the same Leica SP8 and a coverslip corrected HCX IRAPO L 25X/0.95NA water immersion objective (Leica Microsystems). Alexa Fluor 488 and Alexa Fluor 594 were excited with lasers emitting at 488 nm and 552 nm wavelengths respectively. Fluorescence was detected with PMT detectors. SHG signal from nanoKTP in fixed larvae was acquired as in live imaging experiments.

#### Automatic extraction of nanocrystal-labeled vesicle trajectories and transport parameters

Videos were first exported from the Leica LIF proprietary format as stacks of 8-bits TIFF files with a custom FiJi macro relying on the importer function of the Bio-Formats plugin. ^62^ The subsequent data analysis process is automated, using a set of Python functions that we developed and made freely accessible online. ^36^ It consists in two main parts: Part 1 aims at extracting valid trajectories directly from the video files and Part 2 calculates transport parameter observables from the trajectories.

Part 1 is decomposed in three steps: 1) raw trajectories extraction from the videos, 2) filtering directed motions only, 3) regularisation of the localization noise. The stacks are first preprocessed by a white top-hat transform, then processed by the `trackpy`^63^ implementation of the Crocker-Grier algorithm.^64^ This algorithm locates features of a given size (in pixels), brightness (in counts) and minimum separation (in pixels) in each frame (we selected the following parameters of the `batch` function: `diameter=9, minmass=300` and `separation=12`), then links them to form (*x*(*t*), *y*(*t*)) trajectories, as paired arrays of coordinates over time *t*. To this aim we used the `link` function with the following parameters: `search range=6, memory=5, adaptive stop=5 and adaptive step=0.9`. These trajectories are then filtered according to their shape, based on a Mean Square Displacement (MSD) threshold of 300. Trajectories with MSD lower than this threshold were discarded. This filters out non-linear, aberrant trajectories. We found that this algorithm sometimes resulted in a visually singular trajectory being broken down into several smaller ones. We thus rejoined together trajectories of which beginnings and ends are within a given spatiotemporal range from one another. The rejoining thresholds were 10 frames and 40 pixels, *i.e*. the end of one trajectory had to be within 10 frames and 40 pixels distance of the beginning of another one for them to be rejoined. As we aim to measure directed axonal transport, ideal trajectories should progress in a relatively rectilinear motion. Hence, we first rotated each trajectory (*x*(*t*), *y*(*t*)) in a way that the line between the first and furthest points becomes horizontal, and then only selected trajectories which does not deviate from their fit to a third degree polynomial function *f*(*x*) more than 1.4 pixels in standard deviation, *i.e*. trajectories for which ((*y* – *f*(*x*))^2^) < 1.4 pixels, where 〈…〉 is the average on all trajectory points.

The set of selected trajectories was then subjected to a noise filtering, based on a novel algorithm of convex minimization of acceleration, in order to compensate for localization uncertainty. A measured raw trajectory **r**_m_(*t*) extracted from the videos by trackpy consists of the underlying directed motion **r**_d_(*t*) of vesicles moved by the molecular motors, onto which a measurement noise is superimposed that we modelize as a white Gaussian noise **w**_G_(*t*):

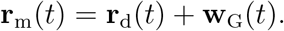

In order to extract the ground truth vesicular motion **r**_d_(*t*) we must correct for the noise first. The vesicular displacements being composed of a succession of phases of almost constant velocity, interrupted by rare pauses with almost zero velocity, the correction implemented relies on the discrimination of these rare but consequent changes in velocity from the permanent ones caused by the noise. An efficient approach to perform this denoising in such a situation is to apply a total variation reconstruction (TVR).^41^ Briefly, we are looking for large changes in the value of the measured (noisy) velocity. Considering 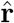 as an estimator of **r**_d_(*t*), our TVR approach consists in minimizing the acceleration 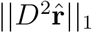 where *D* is the derivation operator relative to the time and 1 stands for the *l*1-norm, under some constraints. The latter is that the estimator 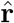 must remain close to the acquired trajectory **r**_*m*_(*t*). With our hypothesis on the noise, we can write that 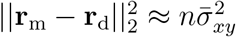, where 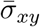 is the average standard deviation of the Gaussian noise **w**_G_(*t*), *n* is the number of frames in the trajectory, and 2 stands for the *l*2-norm.

This leads to the following convex minimization problem:

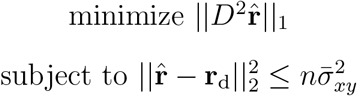

To solve this problem, we use cvxpy (version 1.1.7)^65,66^ Python package that relies on the convex optimization SCS solver^67,68^ (version 2.1.2).

Part 2 consists in parsing the trajectory into Go and Stop phases, corresponding to fast and little to no movement, respectively, using the same confinement ratio (*R*) method used in our previous work.^22^ We determined the threshold *R*_th_ = 0.64 from the minimum of its distribution when considering all trajectories (Supporting Figure S5). At each point *i* we calculate *R_i_* (from 3 preceding and 3 successive positions): *R_i_* > *R*_th_ is associated to a Go phase, while *R_i_* ≤ *R*_th_ is a Stop phase. As the field of view encompasses a very polarized region, with axons projecting into the neuropil, we can discriminate anterograde and retrograde transport based on which direction the particle is moving. Several parameters are then calculated. One general parameter is the *fraction of moving particles*, calculated per field of view as the ratio of the number of trajectories to the number of particle detected in the first frame, and five trajectory-specific parameters:

- **Fraction of moving particles**: ratio of moving particles to non-moving ones. It is calculated by dividing the number of trajectories analyzed for each file (*i.e*. one field of view) by the number of particles found on the first frame of that file. It does not take into account trajectories that were filtered out before analysis, or particles that might appear after the first frame. It is therefore not an absolute measure of the fraction of moving particles, and is only used for relative comparison between experimental conditions.
- **Directionality**: sum of the lengths of the retrograde segments divided by the sum of the absolute values of the lengths of all segments (anterograde and retrograde). A fraction value of 1 correspond to a purely retrograde trajectory, and 0 to a purely anterograde one.
- **Segmental velocity**: speed of the endolysosomal vesicle-embedding particle in μ/s. It is calculated as the average speed of Go phases, which is itself the average of the point velocities for each phase. It is evaluated separately for retrograde and anterograde directions.
- **Pausing frequency**: frequency at which the particle pauses, in number of events per minute. It is calculated as the number of stops divided by the total trajectory time.
- **Pausing duration**: Average time, in seconds, that the particle spends in Stop phases. It is calculated as the average duration of Stop phases in a given trajectory.
- **Run length**: average length, in *μ*m, traveled during Go phases. It is calculated as the average length of GO phases of a trajectory, which is itself the segmental velocity of each phase multiplied by its duration.
- **Theta angle fluctuations** *σ_θ_*: standard deviation of the polar angle *θ* calculated in Go or Stop phases. The angle is first calculated point by point from the intensity (Supporting Data S8, Figure S10), then the standard deviation of that angle for each phase is averaged per trajectory.

#### Statistical analysis and data representation

Statistical differences were analyzed by Wilcoxon signed-rank test (except in Fig. 3, see associated caption). * corresponds to *p* < 0.05, ** *p* < 0.01 and *** *p* < 0.001. All bar plots represent the average of each parameter value, with error bars being ±standard error of the mean (s.e.m.). In all box plots the box delimits the first and third quartiles and the whiskers extend from the first to the ninth decile (10-90%). Outliers are not shown. The median line is surrounded by a notch of which the width defines a confidence interval.

## Acknowledgement

We thank B. Link for sharing the *dync1h1*^mw20^ mutant line. We are grateful to Clément Laigle (Leica Microsystems, France) for his explanation of the conversion of the hybrid detector intensity in photon counts. We thank Eric Larquet for HRTEM acquisition, Carsten Janke for suggesting to use dynapyrazole in replacement of ciliobrevin, Sophie Donnet and Adeline Leclercq Samson for their insight in the statistical analysis of the datasets, Jonathan Piard for the nanoKTP hydrodynamical measurement, Aurélien Ricard for help in data analysis, and Isabelle Garcin for assistance in carrying out first tests in 2D-primary mouse cortical neuron grown on a glass coverslip. Zebrafish experiments were performed at INRAE Infectiology of Fishes and Rodents Facility (IERP-UE907, Jouy-en-Josas Research Center, France doi.org/10.15454/1.5572427140471238E12) which. belongs to the National Distributed Research Infrastructure for the Control of Animal and Zoonotic Emerging Infectious Diseases through *In Vivo* Investigation (EMERG’IN DOI: doi.org/10.15454/90CK-Y371). The authors thank Magali Bouvet and Dimitri Rigaudeau at IERP, for zebrafish husbandry.

This work has received financial support from the CNRS through the MITI interdisciplinary programs (to F.T.), from University Paris-Saclay Strategic Research Initiative “Brainscopes” (French Investments for the Future program grant number 11-IDEX-0003), from ERANET Euronanomed 3 program, through the French National Research Agency (ANR, grant number ANR-18-ENM3-0002 to F.T.), from IHU FOReSIGHT (French Investments for the Future program grant number ANR-18-IAHU-0001 to F.D.B.), from ANR (ANR-20-CE13-0011-02 CodeAX and ANR-19-CE16-0017-01 iReelAx to F.D.B.) and from Fondation pour la Recherche médical (grant #MND202003011485 to F.D.B.). V.B. is funded by a postdoctoral FWO Fellowship (12Y9120N).

## Supporting Information Available

The following files are available free of charge:

- SupportingVideoS1.avi: raster-scanning video from which Fig. 2A was calculated, showing in inverted grey scale the SHG signal of nanoKTP moving in PVN neuron of a zebrafish larva (*kif5aa* wild-type). True frame rate of 20 frames/s, total of 2354 frames. Scale bar: 10 *μ*m.
- SupportingVideoS2.avi: another raster-scanning video showing in inverted grey scale the SHG signal of nanoKTP moving in PVN neuron of a hetegozygous *kif5aa* zebrafish larva. True frame rate of 20 frames/s, total of 2501 frames. Scale bar: 10 *μ*m.
- SupportingVideoS3.avi: example of a raster-scanning video of nanoKTP immobile on a glass coverslip, acquired in the same conditions as the zebrafish larvae, and used to estimate the precision of localization (Supporting Figure 4).

## Supporting information

### 1. Supporting data

#### S1. NanoKTP electron microscopy images and size distribution

**Figure S1:**
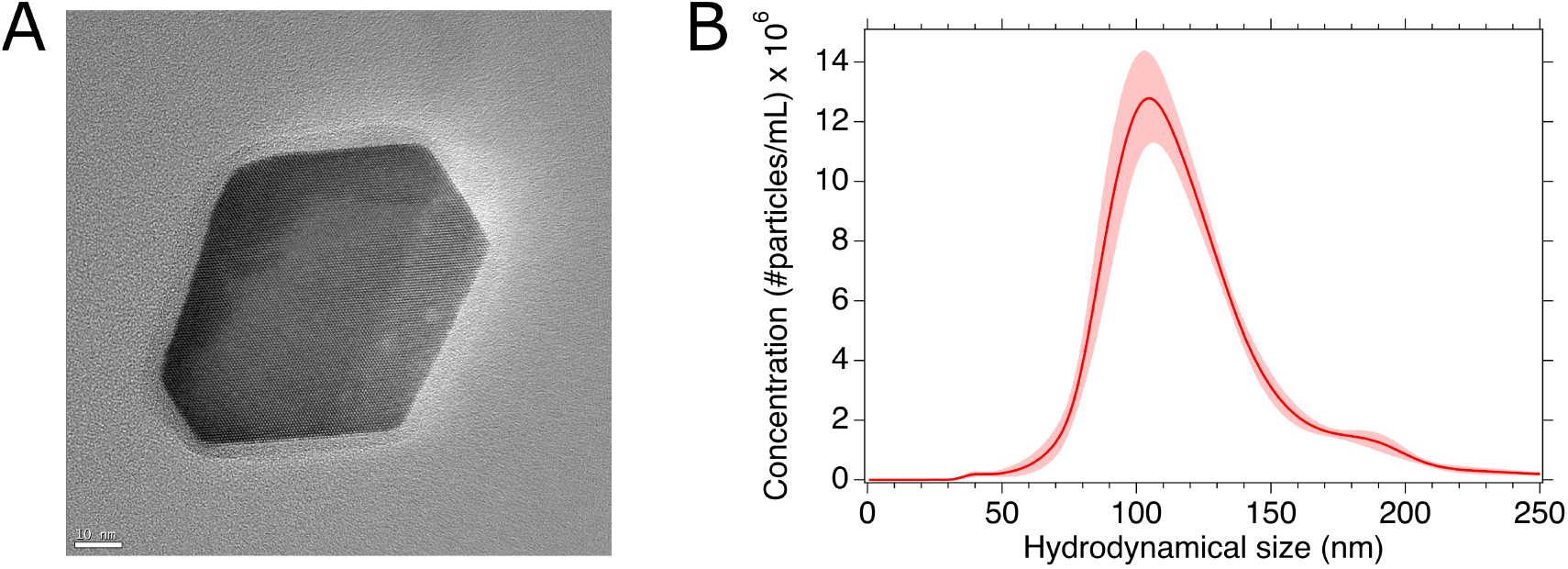
NanoKTP characterization.(A) High resolution transmission electron microscopy image in bright field, acquired with a JEOL JEM-2010F (operated at 200 kV voltage) aberration corrected (Cs: 1.4 mm and Cc: 1.0 mm). The nominal magnification is 400,000 (the effective one on the array detector is 522,049), and the image is acquired −60 nm under-focus. Scale bar: 10 nm. (B) NanoKTP hydrodynamical size distribution as measured by nanoparticle tracking analysis. Red shaded area indicates the limits of five consecutive measurements and the solid line is the average of them.

We used a nanoparticle tracking analysis device (NanoSight NS300, Malvern Panalytical, UK) to measure the hydrodynamical diameter distribution of the size-selected nanoKTP solution (Supporting Figure S1B). We repeated the measurement five times, and the average mean value is 123.8 ± 2.4 nm (standard deviation over the 5 measurements indicated) and an average mode position of 104.0 ± 2.7 nm.

#### S2. Emitted light spectrum

**Figure S2:**
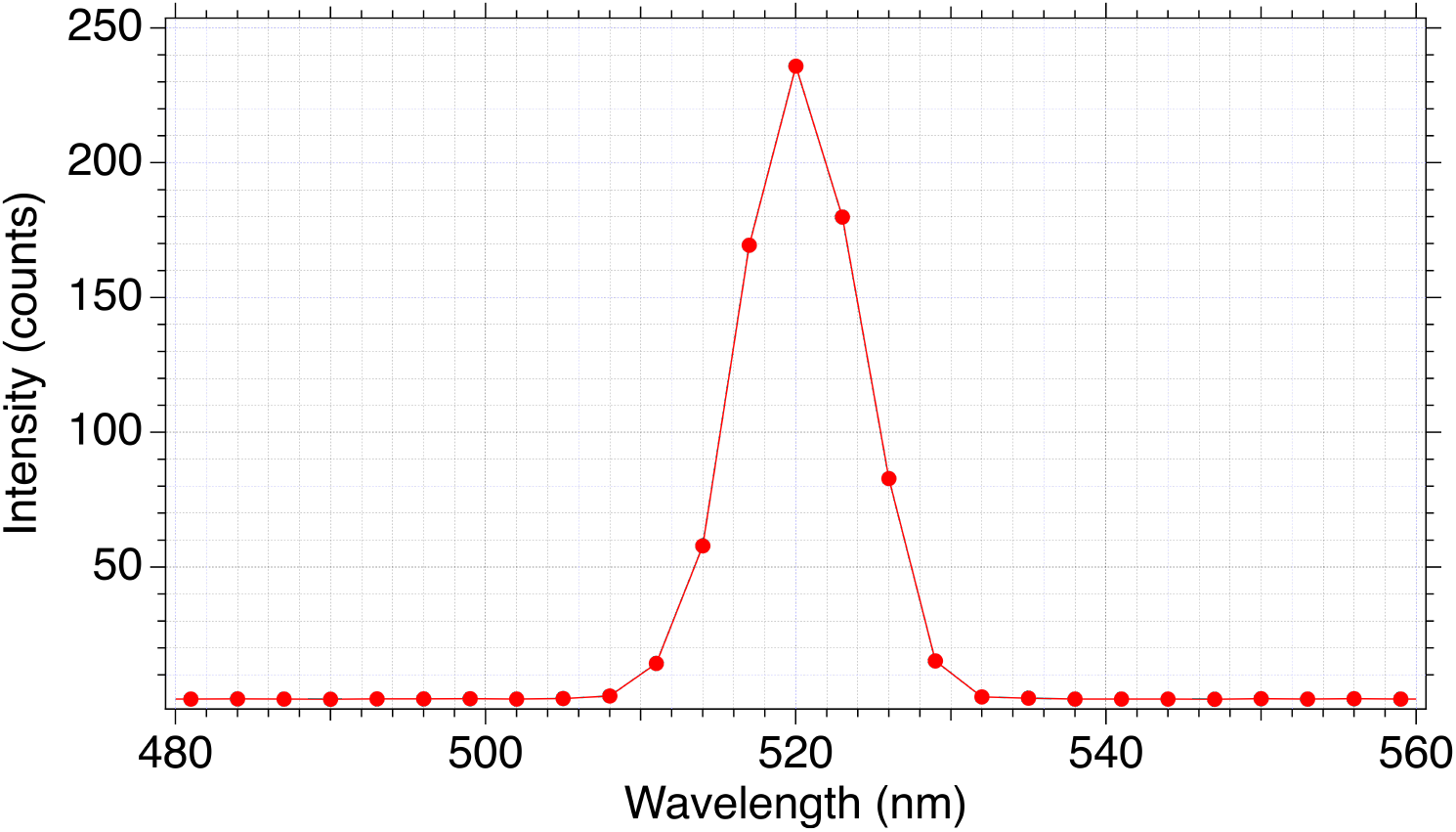
Spectrum of the emitted light from a single nanoKTP as recorded with “lambda scan” mode of the Leica SP8 microscope, showing a narrow band centered on 520 nm corresponding to a SHG signal.

Taking advantage of the prism-based spectrum resolution capability of the Leica SP8 microscope we used, we measured the spectrum of the light emitted from a single nanoKTP excited with the pulsed laser (Chameleon Ultra II, Coherent Inc.) set at the 1040 nm wavelength and observed a peak (Figure S2) which FWHM of 10 nm corresponds to a Fourier transform-limited spectral width of 160 fs. This pulse duration value is in good agreement with the laser specifications of 140 fs, considering the dispersion-induced pulse duration broadening (due in particular to the microscope objective), that is not fully compensated by the pre-compensation module installed between the laser output and the microscope. Moreover, the emission spectrum is perfectly centered on 520 nm, half of the excitation wavelength of 1040 nm. Both the pulse width and duration values confirm that we detect the second-harmonic generation light from nanoKTP.

#### S3. Conversion of 8 bits-encoded hybrid detector intensity levels in photon counts

**Figure S3:**
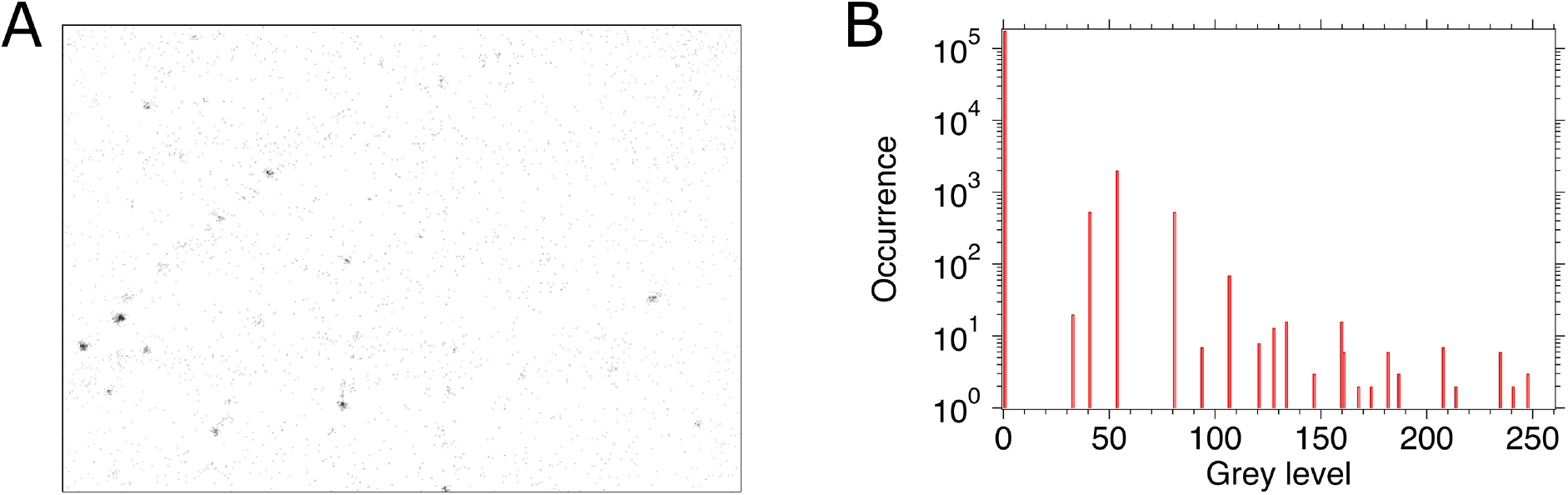
Estimate of the conversion of grey level into numbers of detected photons by the HyD used in the “standard mode”. (A) Frame #665 of Supporting Video S1. (B) Histogram of grey levels present in (A) in logarithmic scale, displaying 21 peaks between the dominant white 0 level (excluded from count) associated to zero detected photons and 255 (black) maximum possible level. The maximum grey level present in (A) is 247.

We estimated the conversion factor between photon counts to 8 bits-encoded intensity grey level from the histogram of grey levels in the whole frame #665 of the Supporting Video S1. Considering that each additional intensity peak results from the counting of one more photon. The maximum intensity peak is the 21^st^ one, and corresponds to a value of 247. Assuming, as a crude approximation, a linear dependence, the photon-to-intensity conversion factor is *K*_conv_ ≈ 247/21 ≈ 11.8 photons per gray level. Let us point out that we have assumed an increment of 1 photon between two consecutive peaks, but it could be larger. Hence, this value of *K*_conv_ is an upper bound. However, the selection of the frame #665 was motivated by two reasons: 1) fact that its largest grey level value of 247 is close to the saturated maximum (255) but lower than it, and 2) because it displays the largest number of peaks of the whole frames. This careful selection should reduce the discrepancy between the estimated conversion factor and the ground truth one.

#### S4. Precision of localization of the nanoKTP: theoretical estimate and comparison with its measurement

**Figure S4:**
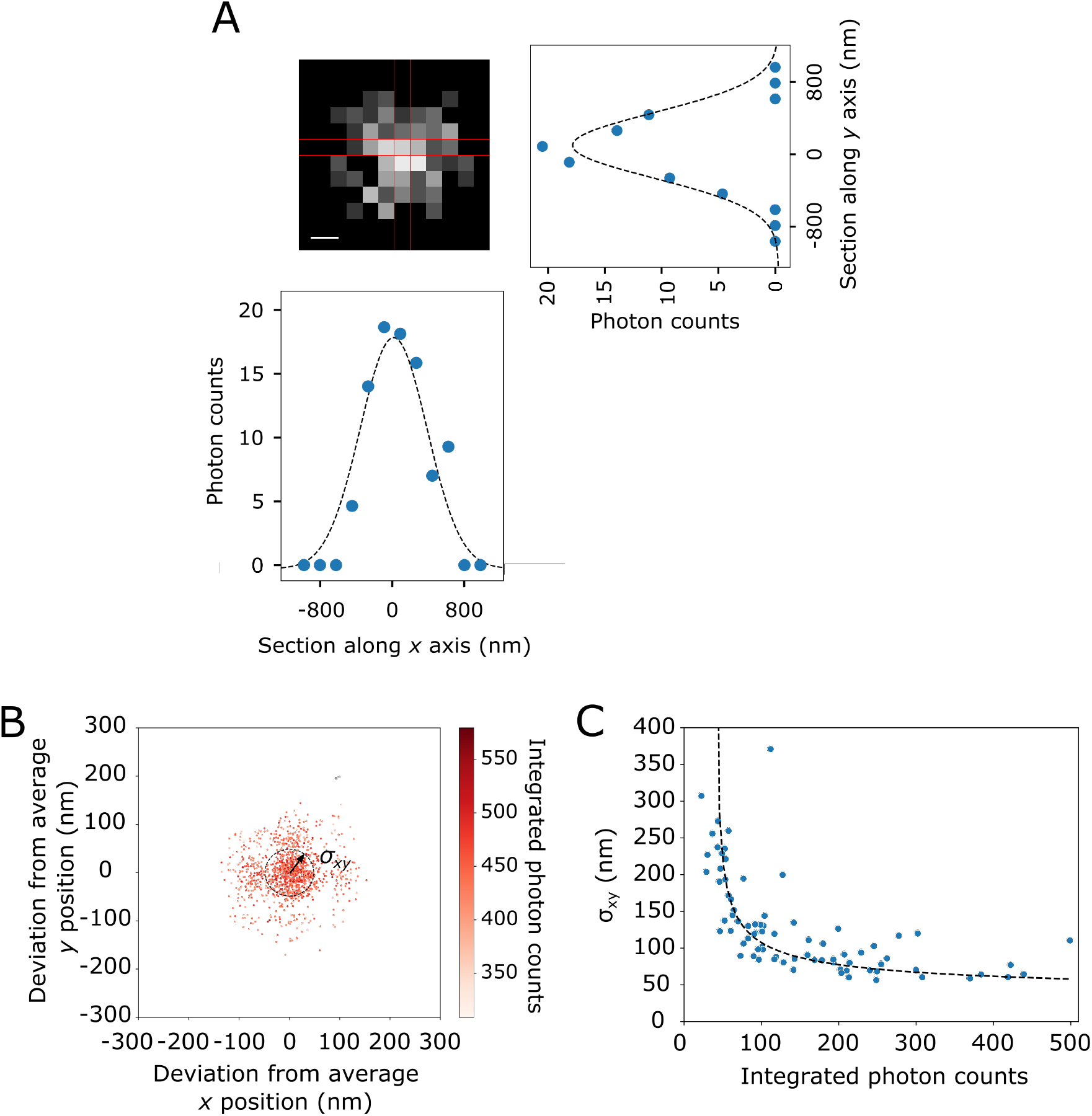
Measurement of the imaging precision of localization. (A) SHG spot from one nanoKTP immobilized on a glass coverslip. Bottom and right are intensity sections along lines *y* = 0 and *x* = *O* respectively, blue points being the experimental values and the dashes black lines being Gaussian fits. (B) Scatter plot of Gaussian fit center position of the same spot than in (A) in consecutive frames of Supporting Video S3. *σ_xy_* arrow and dashed line circle represents the standard deviation of the distribution of positions centered on the average one. (C) Distribution of *σ_xy_* for an ensemble of SHG spots of different mean integrated photon counts. The dashed black line is an exponential fit.

We determined the theoretical precision of localization of a single nanoKTP from the number of photons detected during the raster scan of its SHG emission. This limit is defined as the Cramèr-Rao bound (CRB) (*σ*_CRB_, being the standard deviation) from Fisher information theory. As the photons constituting SHG spot are collected pixel after pixel along the raster scan and not at once on the pixels of an array detector (as in Single Molecule Localization Microscopies, SMLM), different theoretical limits can be achieved for the two acquisition methods^1^. However, it can be shown in the case where we only take into account the photon shot noise^2^ that for a Gaussian shape excitation beam (which is a very good approximation of the focused excitation laser spot), both the SMLM and raster-scanning imaging modes yield almost the same lower bound^3^: 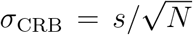, where s is the PSF standard deviation, and *N* the total number of photons detected over the spot.

We estimated s from one of the brightest SHG spots associated to a nanoKTP immobilized on a glass microscope coverslip. We extracted such spot, displayed on Supporting Figure S4A, from Supporting Video S3 which is one of the seven one-minute duration videos of SHG signal from immobile nanoKTP we recorded. As the nanoparticle have a non negligible size relative to the expected s value, the SHG spot intensity distribution results from the convolution of the particle profile (approximated to a sphere of diameter *d*^nanoKTP^ = 120 nm) and the PSF_exp_ function. Figure S4A shows one of the brightest spots and Gaussian fits of two orthogonal centered cross-sections with standard deviations 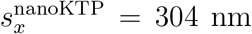 and 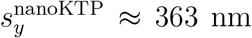. As a crude approximation: 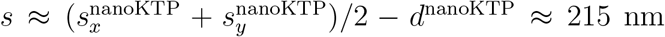. By integrating the grey levels of intensity of all pixel composing Supporting Figure S4A spot, and using the conversion factor determined in section, we obtain *N*_avg_ = 439, leading to 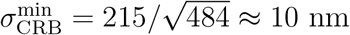.

The experimental value we measured for this spot is the standard deviation of the Gaussian fit center positions in consecutive frame of the video (Supporting Figure S4B), which is *σ_xy_* ≈ 50 nm. This value appears to be larger than the theoretical estimate 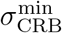, probably due to mechanical vibrations. We then considered other spots of different intensities from all the seven videos. Figure S4C displays the variation of *σ_xy_* with the integrated photon counts per spot.

#### S5. Confinement ratio threshold to parse the trajectory in Go and Stop phases

**Figure S5:**
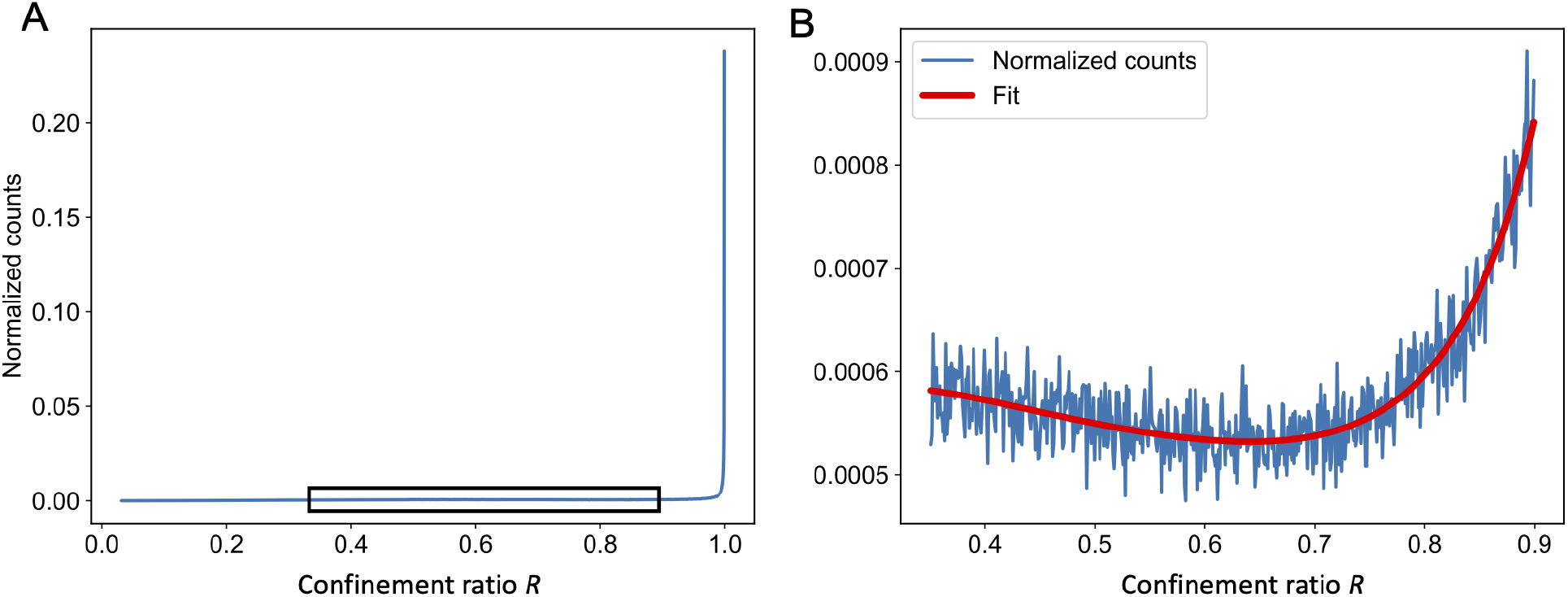
Determination of the confinement ratio threshold *R*_th_ to parse the trajectories in Go and Stop phases of motion. (A) Histogram of the confinement ratio values for all trajectories of all conditions of *kif5aa* larvae. The black rectangle defines the region where we can observe a minimum of *R*. (B) Zoom on the black rectangle region: the minimum *R*_th_ = 0.64 is obtained from a polynomial fit (red line) of the histogram (blue line).

The confinement ratio *R_i_* is determined at each point i along all the trajectories over a duration of 6Δ*t* = 0.3 s (Δ*t* = 50 ms being the duration of one full frame scan), corresponding to 3 preceding plus 3 successive measured positions of the nanoparticle. A unit value corresponds to a perfectly directional motion, whereas a zero value means that the particle came back at the exact same location after a duration of 6Δ*t*. The histogram of {*R_i_*} is shown on Supporting Figure 5. To analyze the intraneuronal transport from our experiment, we separate the globally directional motions from the more diffusive ones, characterized by lower confinement ratio values. We find a threshold between these two behaviors by representing the logarithmic counts as a function of *R* and isolating a local minimum using a 5^th^-order polynomial fit. The minimum is found for *R*_th_ ≈ 0.64, which is the threshold used to parse each trajectory in Stop and Go phases.

#### S6. Absence of effect of nanoKTP on lysosomes axonal transport parameters

To evaluate the potential impact of nanoKTP injection on axonal transport, we labeled lysosomes with LysoTracker live dye, traced their motion at the same frame rate (20 fps) than nanoKTP and extracted their transport parameters with the same analysis pipeline as the one used for nanoKTP. The extraction of trajectories was not as efficient for LysoTracker positive vesicle than for nanoKTP, due to a lower signal-to-background ratio, as can be seen in Fig.S7. However, we collected enough data in the different conditions and Figure S6 shows that lysosomes (LysoTracker-labeled vesicles) axonal transport parameters are not altered by nanoKTP injection.

#### S7. Colocalization of nanoKTP with lysosomes

Considering that we inject the nanoKTP 24h before imaging, we anticipate that after their endocytosis, they are most likely located in late endosomes and lysosomes. To check this hypothesis, we conducted an additional experiment in which the nanoKTP injected larvae were bathed with LysoTracker Red live imaging dye, and then imaged in two-color channels (green for nanoKTP, red for LysoTracker) at 20 fps. We only focused on mobile nanoKTP (with directed motion), which are the only one for which we can ascertain the uptake. Figure S7(A-C) displays an example of a field-of-view with two mobile nanoKTP colocalizing with LysoTracker Red vesicles. Fig.S7D is the statistical summary, concluding to about 56% of the nanoKTP in late endosomes or lysosomes.

**Figure S6:**
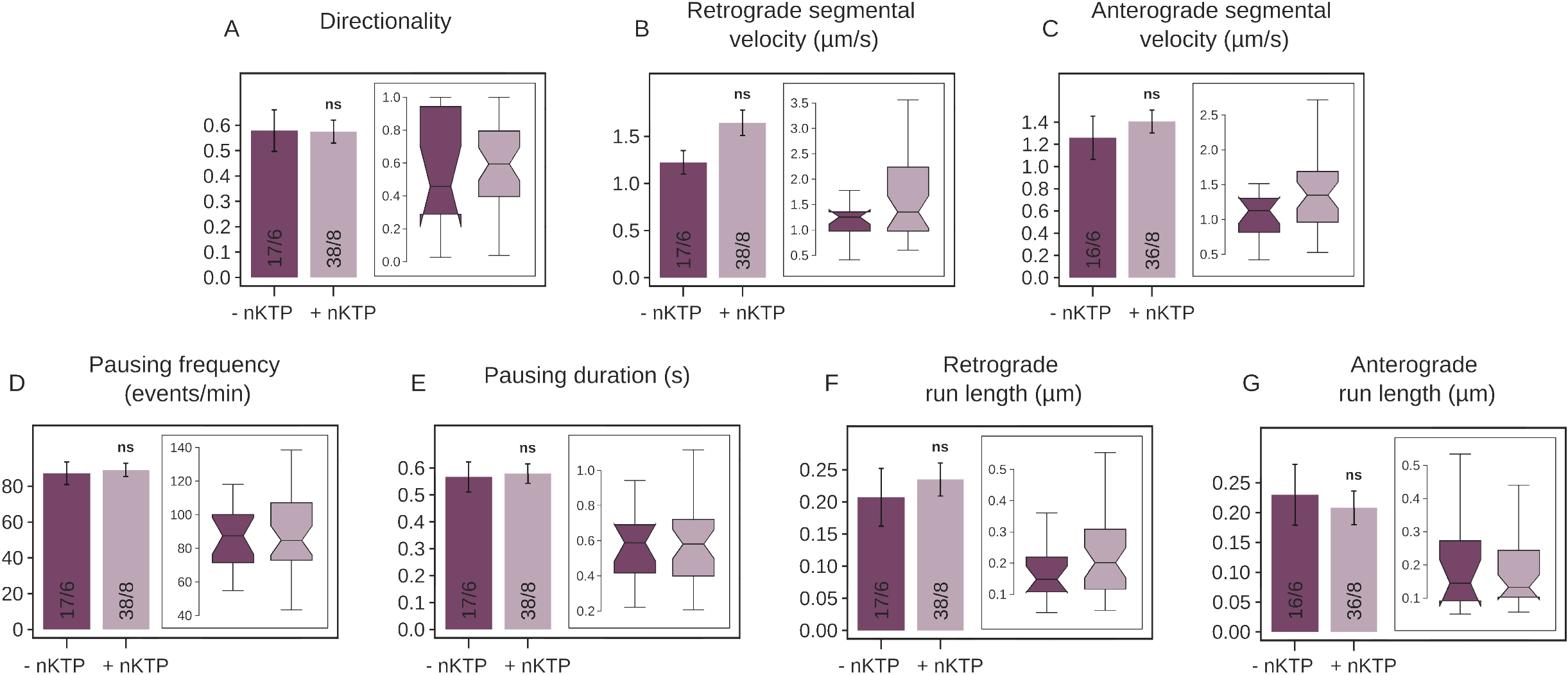
Effects of nanoKTP on lysosome axonal transport parameters. We bathed Zf larvae at 4 dpf with LysoTracker Red to trace lysosomes and compared two conditions: with nanoKTP (+nKTP) injected at 3 dpf or without (-nKTP). Lysosome transport parameters were extracted with the same analysis pipeline as nanoKTP. As a reminder, the directionality (A) is the cumulative length of retrograde segment divided by the total length of the trajectory. (B) and (C) are the retrograde and anterograde segmental velocities, respectively. (D) and (E) are the pausing frequency and pausing duration respectively, while (F) and (G) are the anterograde and retrograde runlengths, respectively. None of these parameters present significant differences between +nKTP and -nKTP conditions (from (A) to (G): *p* = 0.87, 0.15, 0.18, 0.78, 0.85, 0.31 and 0.97). Numbers in the bars are # trajectories/# of animals.

**Figure S7:**
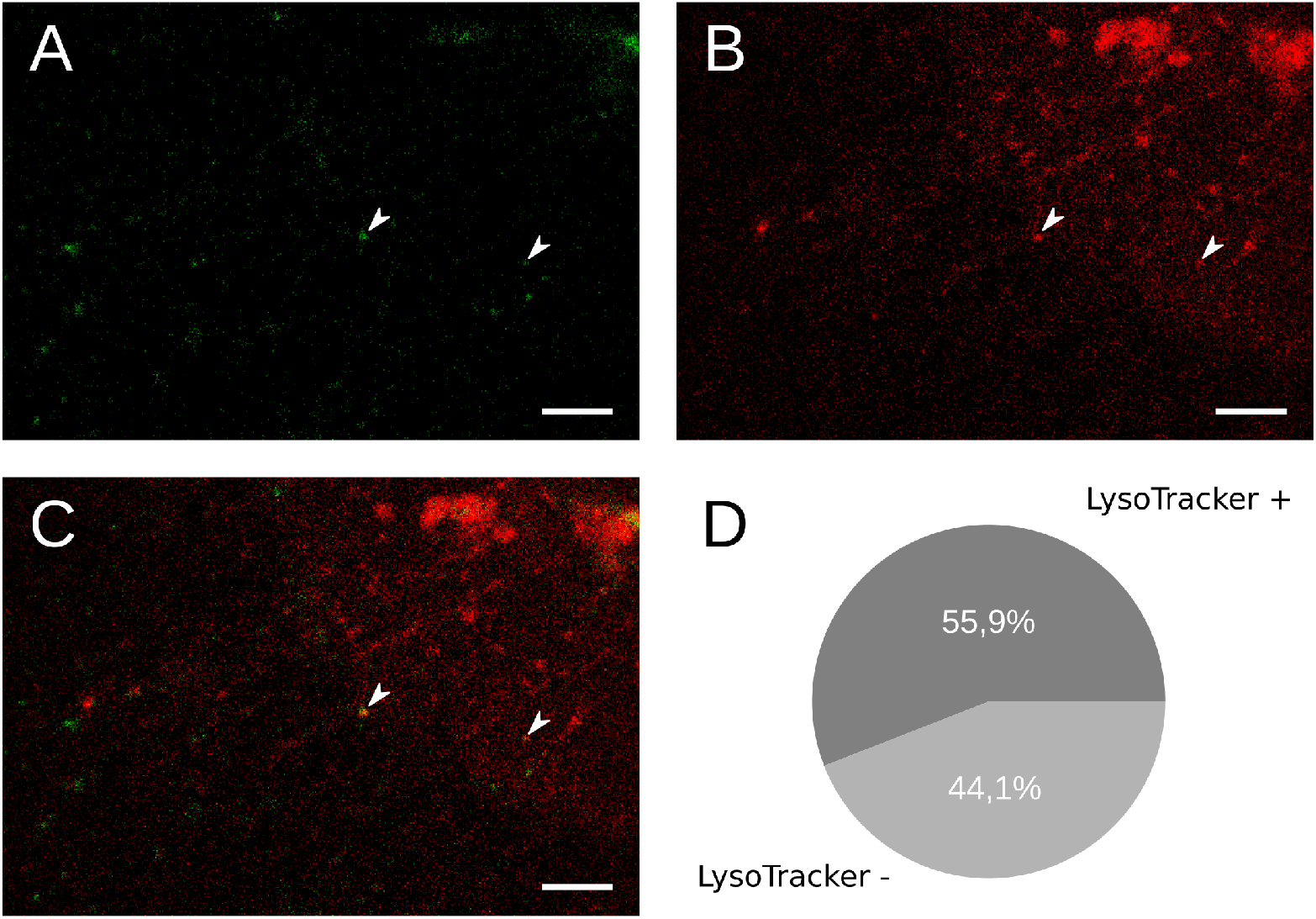
Colocalization of mobile nanoKTP with lysosomes. (A) One frame of the green channel (nanoKTP) of a two-color (green and red, see Materials and Methods) video. (B) LysoTracker Red channel of the exact same frame. (C) Merge of green and red channels. The two arrow heads point towards nanoKTP having directed motions and colocalizing with LysoTracker Red positive compartment. (D) Result of the mobile nanoKTP-LysoTracker colocalization analysis from one experiment carried out for on 8 larvae, 62 fields of views (from both brain sides), leading to a total of 38 nanoKTP trajectories. LysoTracker +/- correspond respectively to nanoKTP containing vesicles that are positive/negative to LysoTracker signal. Scale bars: 10 μm.

#### S8. Reversal of directionality quantification for dynapyrazole-treated larvae and *kif5aa* larvae

The directional switch frequency, displayed on Figure S8 in the case of dynapyrazole treatment and for *kif5aa* larvae, is calculated as the number of events where a Go phase of a given direction is followed by a Go phase of the opposite direction, after a Stop. The number of events was then normalized by the duration in seconds of the trajectory.

**Figure S8:**
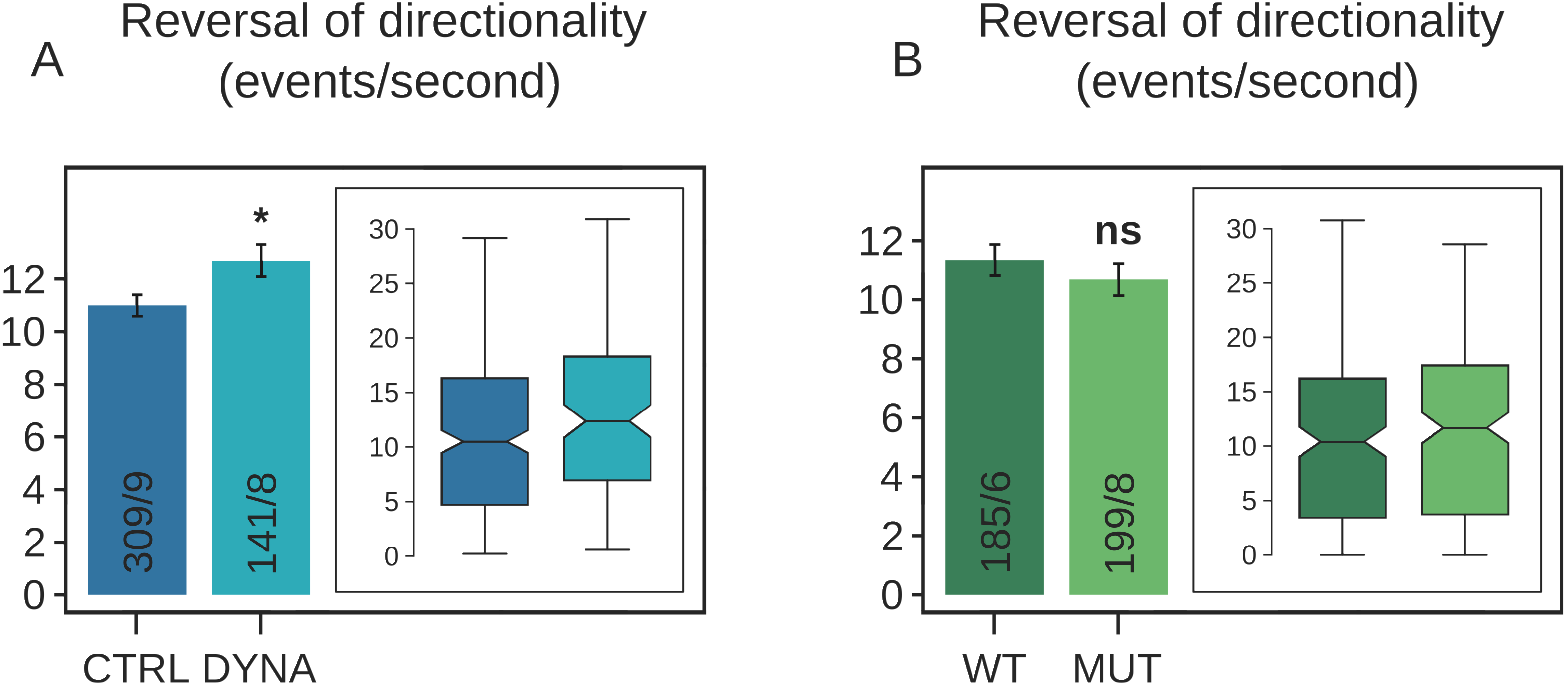
Directional switch frequency in the case of dynapyrazole treatment (A) and for *kif5aa* larvae (B). Dynapyrazole increases the switch frequency from to (*p* = 0.025). There is no switch frequency difference between WT and MUT *kif5aa* (*p* = 0.352)). Numbers in the bars are # trajectories/# of animals

#### S9. Separate quantifications of pausing durations for pauses between two anterograde or two retrograde phase of motion, or between phase of opposite directions

The separate quantification of pausing duration displayed in Figure S9 excludes, by construction, pauses at the extremities of trajectories.

#### S10. Collection efficiency of SHG light emitted by a single monocrystalline nanoKTP

The geometry and notations are defined in the Supporting Figure S10A.

As *d*_33_ = 16.9 pm/V nonlinear coefficient of KTP dominates the others (ranging from 1.9 to 4.3 pm/V), we consider the SHG emission as solely due to an induced dipole **p**_3_ along the nanocrystal *c*-axis **u**_3_:

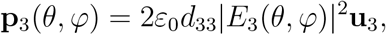

where *E*_3_(*θ, φ*) is the projection of the circularly polarized excitation laser electric field **E**^inc^ on **u**_3_. In the following we will consider *φ* = 0 (**e**_x_ along the direction of the projection of **p**_3_ in the sample plane (Oxy)), without any restriction of generality due to the cylindrical symmetry along the microscope objective axis (O*z*), so that **u**_3_ = sin *θ***e**_*x*_ + cos *θe_z_*, with *θ* being the polar orientation angle of **p**_3_ relative to (O*z*)

**Figure S9:**
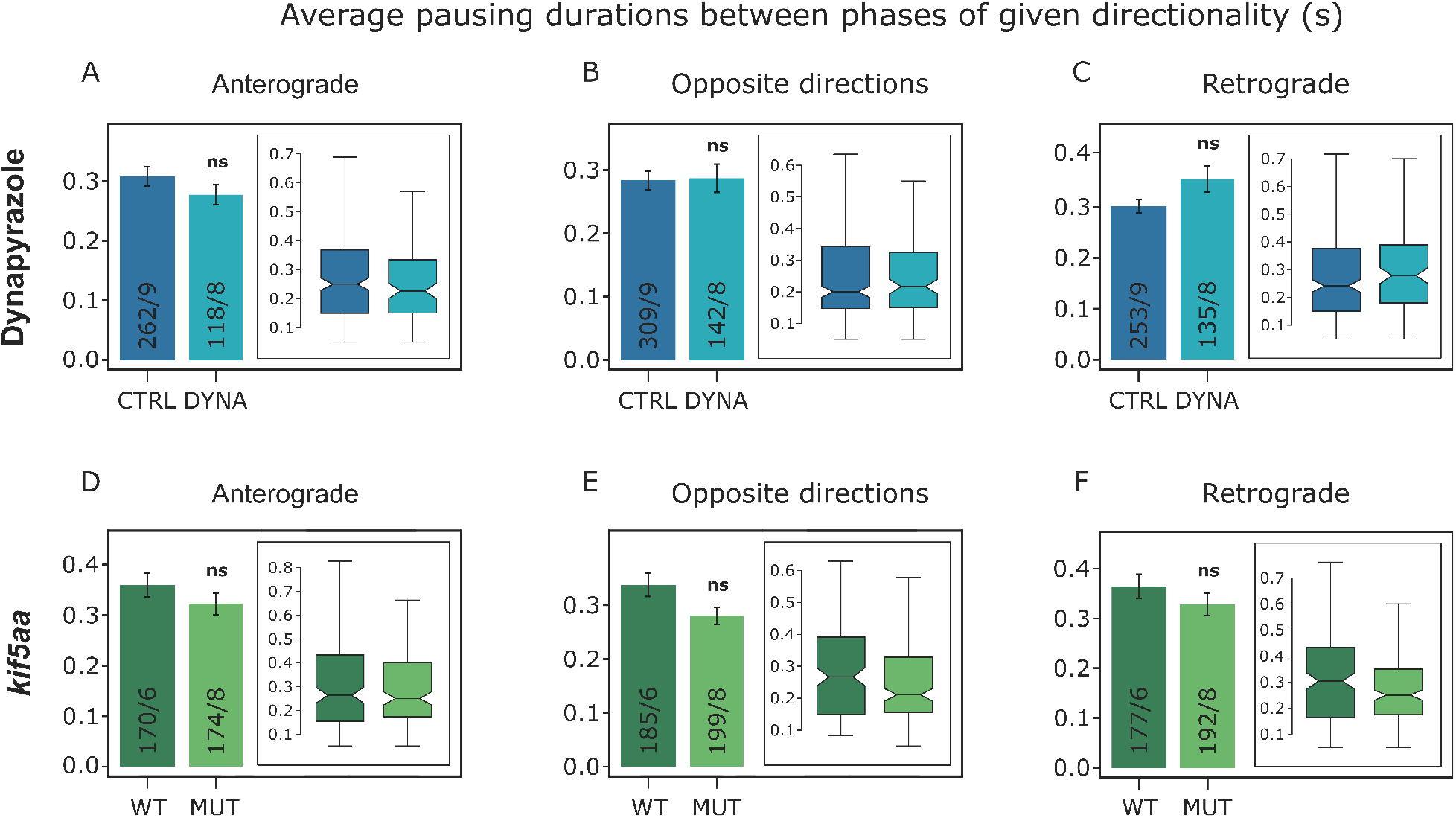
Average pausing durations between phases of motion of given directionality, in the case of DYNA (A-C) and *kif5aa* (D-F) experiments. Pausing duration between two anterograde phases (A,D), two retrograde phases (C,F) or two phases of opposite direction of motion (B,E). p-values are, from A to F, respectively: 0.57, 0.67, 0.09, 0.58, 0.16 and 0.21. Numbers in the bars are # trajectories/# of animals.

The SHG total collected power *P*_2*ω*_(*θ*) (where *ω* is the pulsation of the excitation laser) results from the angular integration of all Poynting vectors of electromagnetic fields at 2*ω* pulsation, with unitary vector directions **u** = sin *θ′* cos *φ′***e**_x_ + sin *θ′* sin *φ′***e_y_** + cos *θ′***e_z_**.

We first consider the radiated power per unit solid angle of collection around **u**, which is known to have the form:

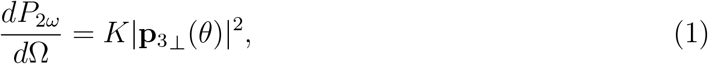

where *K* is a constant, and **p**_3⊥_ ≡ **p**_3_ – (**p**_3_.**u**)**u** is the the projection of **p**_3_ in the plane perpendicular to **u**.

**Figure S10:**
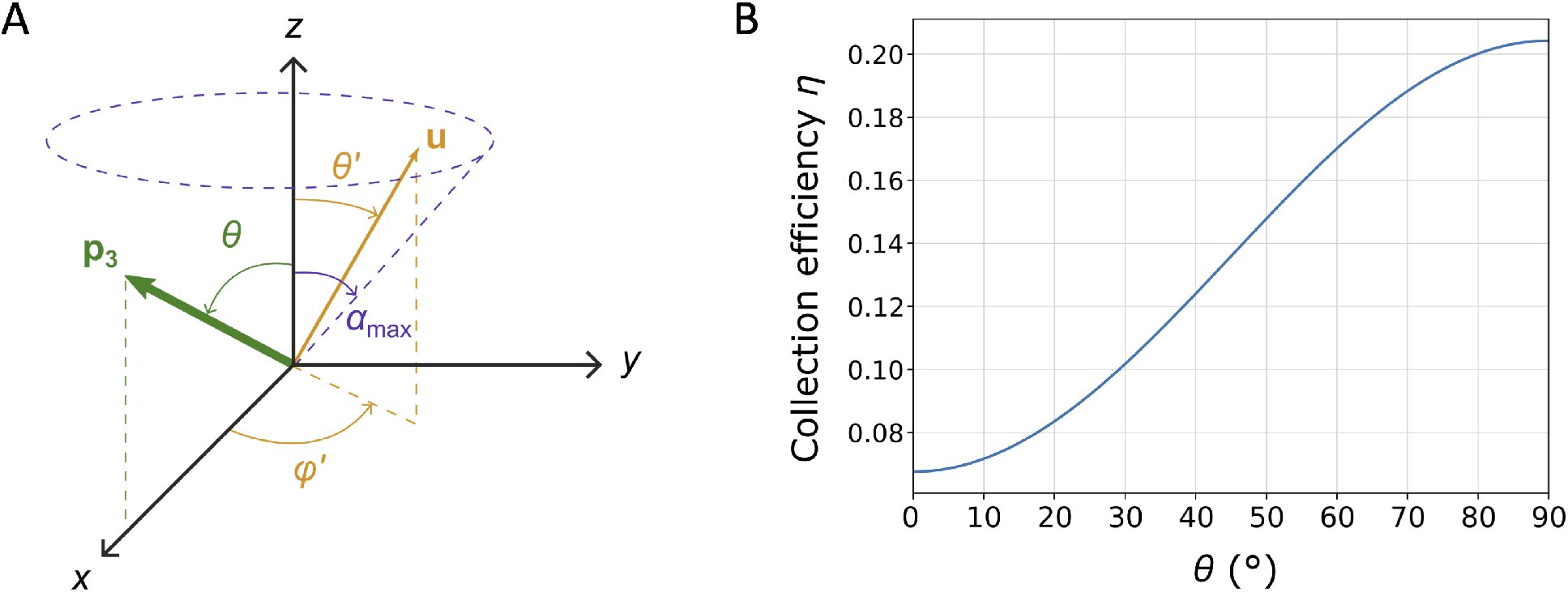
Collection efficiency of the SHG signal from a single nanoKTP. (A) Geometry for the calculation of the collection efficiency. The SHG is excited with a pulsed laser beam focused in the sample plane orthogonal to the microscope objective symmetry axis (O*z*). The excitation light is circularly polarized to ensure equal excitation of nanoKTP with different crystalline axis orientations. We model the nanocrystal second-order nonlinear response as the one of a dipole **p**_3_ (along the crystal *c*-axis). For a given nanocrystal, we can select the (O*x*) axis to match the projection of this dipole in the sample plane, with no loss of generality. SHG emission is collected through the same objective. **u** denotes one unitary vector direction of SHG wave vector, with dircetion defined by *φ′* and *θ′* angles. *θ′* ranges between 0 and *α*_max_, defined through *NA* = *n*_w_ sin *α*_max_, with *NA* = 0.95, the numerical aperture of the microscope objective, and *n*_w_ = 1.3 the index of refraction of the aqueous imaging medium. In our case *α*_max_ ≈ 50.3°. (B) SHG collection efficiency *η*(*θ*) of a nanoKTP as a function of its polar orientation *θ*, with respect to the microscope objective axis (O*z*). *η*(*θ*) varies as sin^2^ *θ* with a contrast depending on the numerical aperture NA. Plot drawn for water immersion microscope objective with *NA* = 0.95.

One can show that |**p**_3⊥_|^2^ = |**p**_3_1^2^*M*(*θ, θ′, φ′*), where:

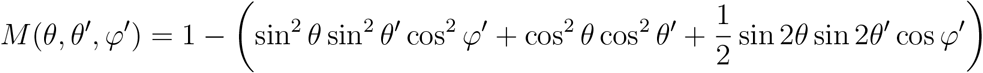

Hence, *P*_2*ω*_ (*θ*) is obtained by integrating equation (1) over the solid angles *d*Ω, with the upper limit *α*_max_ of integration on *θ′* defined by the numerical aperture *NA* of the microscope objective (*NA* = *n*_w_ sin *α*_max_, where *n*_w_ = 1.3 is the index of refraction of water):

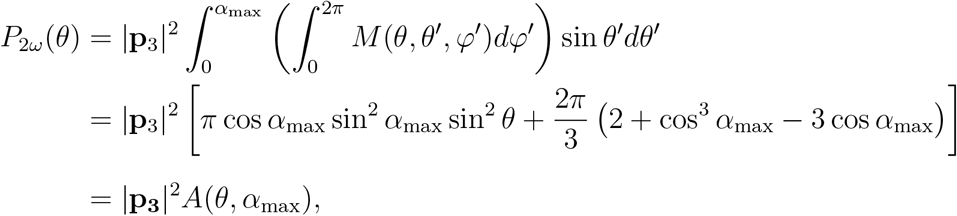

where the notation *A*(*θ, α*_max_) ≡ *B*(*α*_max_)sin^2^ *θ* + *C*(*α*_max_) is introduced to clarify the expression.

Of note, integrating on 4*π* steradian results in 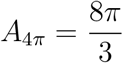. The power *P*_2*ω*_(*θ*) can thus be normalized by this value, leading to the collection efficiency *η*(*θ*):

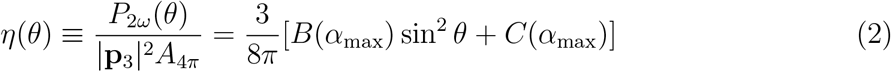

Supporting Figure S10B displays *η*(*θ*).

#### S11. Example of *θ*(*t*) retrieval from SHG intensity variation. Case of static nanoKTP

Supporting Figure S6 shows an example of *θ*(*t*) retrieval from a nanoKTP with a directed motion. We assumed that the detected SHG intensity (in unit of extrapolated photon counts per frame duration) 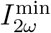 and 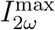 are reached for *θ* = 0° and 90° respectively, so that *θ*(*t*) is retrieved, in radians, by:

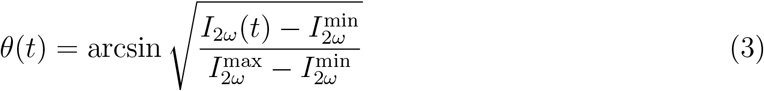

In order to reduce the sensitivity of *θ*(*t*) retrieval to SHG intensity large but short fluctuations, we applied a Savitzky-Golay filter to *I*_2*ω*_(*t*) of all trajectories (orange curve in Fig.S11A) prior to the extraction of *θ*(*t*) (Fig.S11B) by equation 3. Fig.S11C displays the SHG intensity time trace of one of the immobilized nanoKTP used to determined the localization precision (Fig.S4). As our method to retrieve *θ* angular variations cannot be applied to a static particle, we cannot convert the intensity fluctuation into angular fluctuations for static particles. However, we believe that *I*_2*ω*_(*t*) fluctuations mainly result from mechanical vibrations resulting in slight change of focus and excitation intensity in the sample plane. Indeed, the average intensity and standard deviation for the static particle considered are respectively 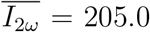 and *σ_I_* = 26.8 counts (per time frame), while the shot-noise limit for 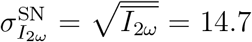. The excess of noise is consitent with our localisation precision not limited by the shot-noise, but rather by mechanical vibrations (Fig.S4C).

**Figure S11:**
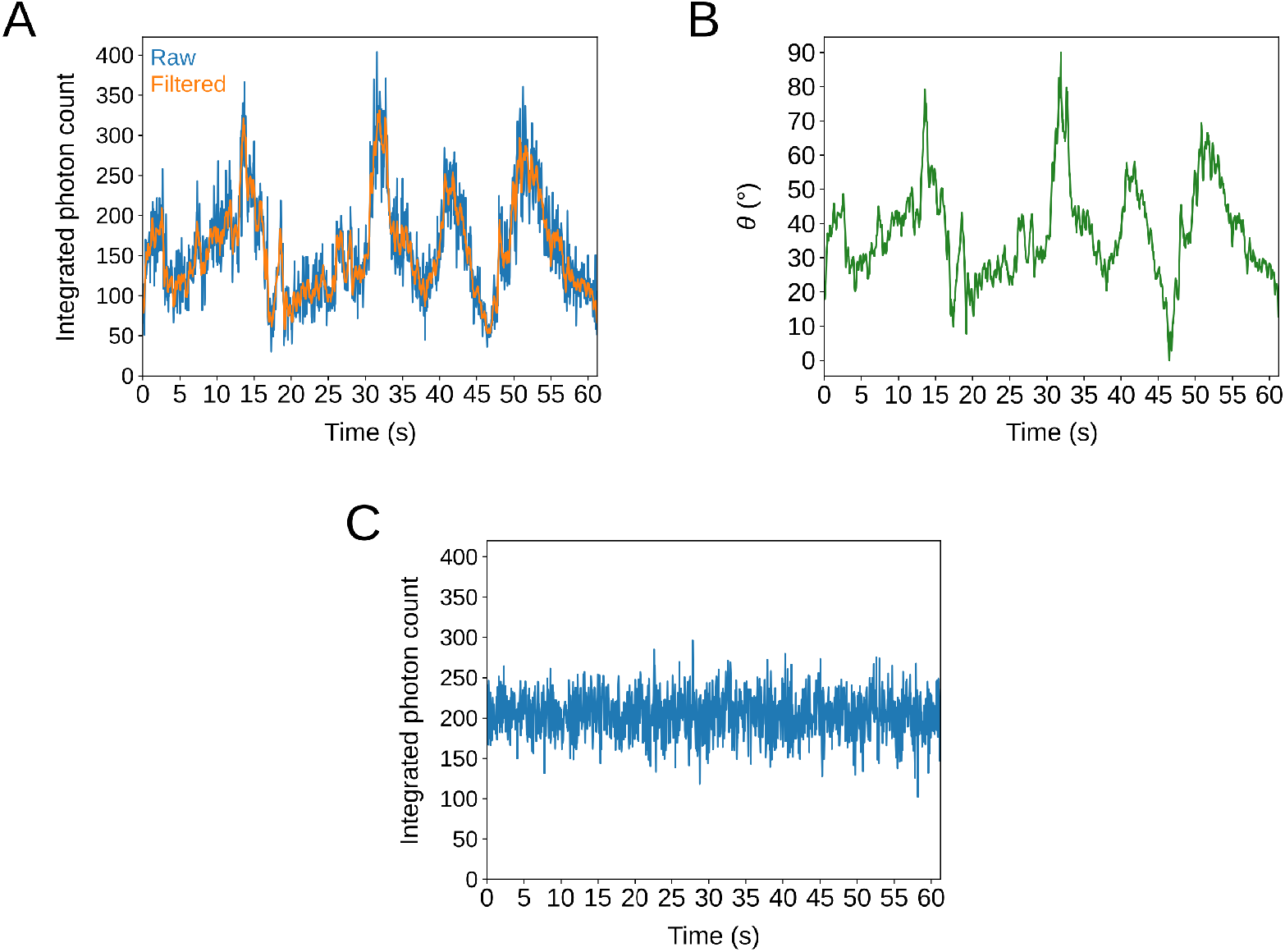
SHG detected intensity *I*_2*ω*_(*t*) variation of one mobile and one static nanoKTP, and *θ*(*t*) inference. (A) SHG intensity time trace from a nanoKTP following a directed motion (inset) in an axon of HOM *kif5aa* larva. The blue trace is the raw data with 50 ms time interval between consecutive frames. The orange superimposed trace results from filtering the raw trace with a Savitzky-Golay filter using a sliding window of 11 time intervals and a third order polynomial fit. (B) *θ*(*t*) time trace inferred from (A) *I*_2*ω*_(*t*) using equation (3), assuming that *I*_2*ω*_ minimum and maximum correspond respectively to 0° and 90°. (C) Intensity fluctuations of a static nanoKTP immobilized on a glass microscope coverslip. Intensity is expressed in photon counts as estimated from the conversion detailed in Supporting Data S3.

#### S12. Fluctuations of orientations of nanoKTP-labeled endolysosomal vesicle in dynapyrazole-treated zebrafish larvae

**Figure S12:**
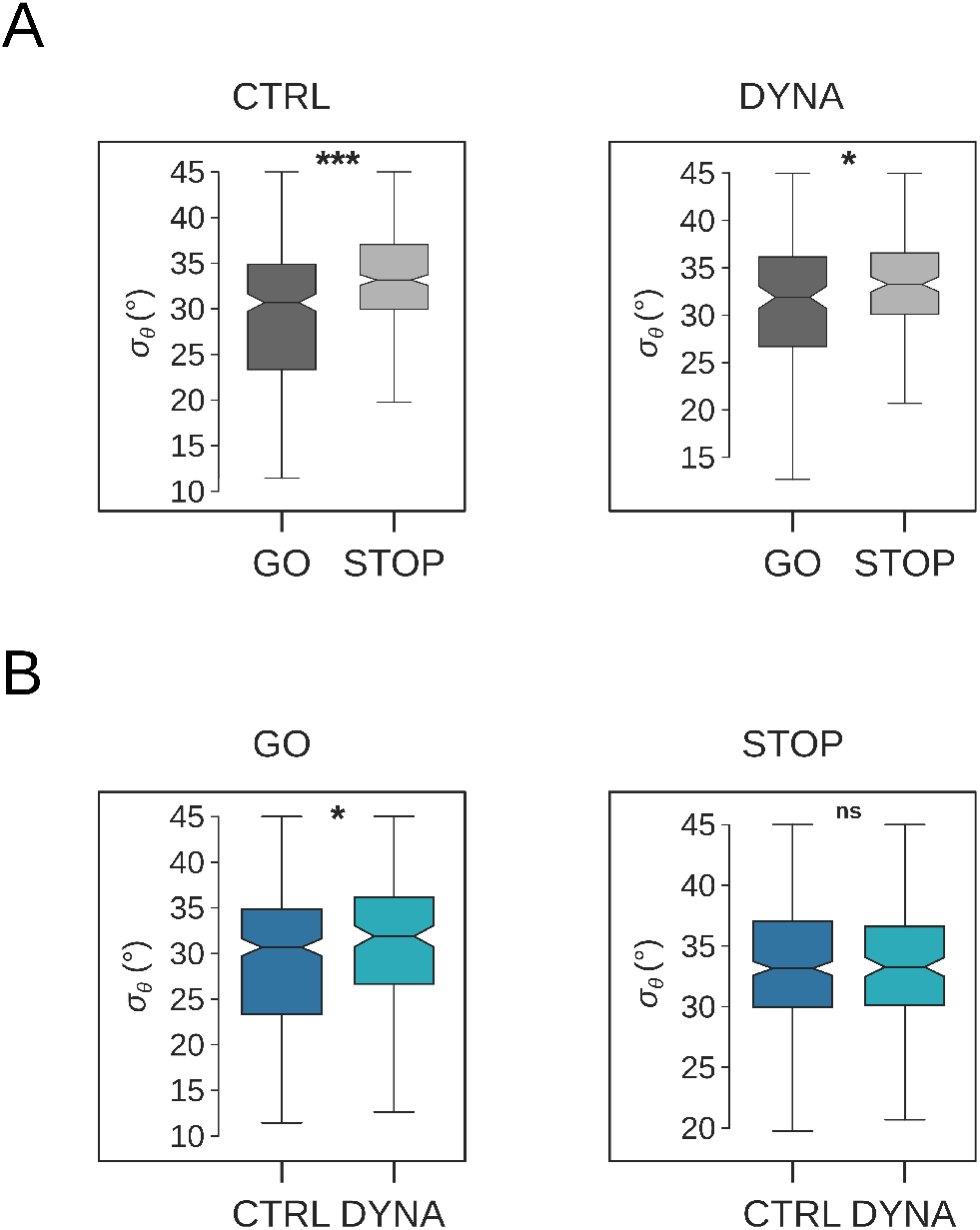
Fluctuation of orientation *σ_θ_* of nanoKTP-labeled vesicle during the Go and Stop phases of motion in neurons of the brain of Zf larvae treated with dynapyrazole-A, compared to controlled untreated ones. As reported for *kif5aa* larvae (Figure 6), *σ_θ_* is larger in the Stop phases of motion than in the Go ones for both Control (increase of 8%) and DYNA conditions (increase of 8.0%), *p* = 2.2 × 10^-12^ and 0.02 respectively. Comparing *σ_θ_* between different conditions for each phase of motion, we observe that DYNA treatment induces an increase of 4.8%, from 29.3° to 30.7° in average, of *σ_θ_* in the case of the Go phases (*p* = 0.03) but not in the Stop phases (*p* = 0.44).

Supporting Figure S12 displays the standard deviation *σ_θ_* quantifying the fluctuation of polar angle *θ* for larvae treated by DYNA compared to untreated control, in Go and Stop phases.

### 2. Supporting videos

Supporting Video S1 is the raster-scanning video from which Fig. 2A was calculated, showing in inverted grey scale the SHG signal of nanoKTP moving in PVN neuron of a zebrafish larva (*kif5aa* wild-type). True frame rate of 20 frames/s, total of 2354 frames. Scale bar: 10 μm. Supporting Video S2 is another raster-scanning video showing in inverted grey scale the SHG signal of nanoKTP moving in PVN neuron of a hetegozygous *kif5aa* zebrafish larva. True frame rate of 20 frames/s, total of 2501 frames. Scale bar: 10 *μ*m. Supporting Video S3 displays SHG of individual nanoKTP drop-casted on a glass coverslip, that was used to infer the precision of localization displayed on Supporting Figure S4, with 10 other similar videos. They all were acquired in the exact same conditions as the videos of nanoKTP directed motion in the ZF larvae brain, *i.e*. in particular at 20 frame/s, inside the thermally controlled cage incubator.

## References

1. Millecamps, S.; Julien, J.-P. Axonal transport deficits and neurodegenerative diseases. Nature Reviews Neuroscience 2013, 14, 161–176.

2. Guedes-Dias, P.; Holzbaur, E. L. F. Axonal transport: Driving synaptic function. Science 2019, 366, eaaw9997.

3. Maday, S.; Twelvetrees, A.; Moughamian, A.; Holzbaur, E. Axonal Transport: Cargo-Specific Mechanisms of Motility and Regulation. Neuron 2014, 84.

4. Hirokawa, N.; Noda, Y.; Tanaka, Y.; Niwa, S. Kinesin superfamily motor proteins and intracellular transport. Nature Reviews Molecular Cell Biology 2009, 10, 682–696.

5. Canty, J. T.; Tan, R.; Kusakci, E.; Fernandes, J.; Yildiz, A. Structure and Mechanics of Dynein Motors. Annual Review of Biophysics 2021, 50, 549–574.

6. Surana, S.; Villarroel-Campos, D.; Lazo, O. M.; Moretto, E.; Tosolini, A. P.; Rhymes, E. R.; Richter, S.; Sleigh, J. N.; Schiavo, G. The evolution of the axonal transport toolkit. Traffic 2020, 21, 13–33.

7. Bilsland, L. G.; Sahai, E.; Kelly, G.; Golding, M.; Greensmith, L.; Schiavo, G. Deficits in axonal transport precede ALS symptoms in vivo. Proceedings of the National Academy of Sciences 2010, 107, 20523–20528.

8. Sleigh, J. N.; Tosolini, A. P.; Schiavo, G. Axon Degeneration, Methods and Protocols. Methods in molecular biology (Clifton, N.J.) 2020, 2143, 271–292.

9. Bartheld, C. S. v. Axonal transport and neuronal transcytosis of trophic factors, tracers, and pathogens. Journal of Neurobiology 2004, 58, 295–314.

10. Sleigh, J. N.; Vagnoni, A.; Twelvetrees, A. E.; Schiavo, G. Methodological advances in imaging intravital axonal transport. F1000Research 2017, 6.

11. Mandal, A.; Pinter, K.; Drerup, C. M. Analyzing Neuronal Mitochondria in vivo Using Fluorescent Reporters in Zebrafish. Frontiers in Cell and Developmental Biology 2018, 6, 144.

12. Bercier, V.; Hubbard, J. M.; Fidelin, K.; Duroure, K.; Auer, T. O.; Revenu, C.; Wyart, C.; Bene, F. D. Dynactin1 depletion leads to neuromuscular synapse instability and functional abnormalities. Molecular Neurodegeneration 2019, 14, 27.

13. Plucinska, G.; Paquet, D.; Hruscha, A.; Godinho, L.; Haass, C.; Schmid, B.; Misgeld, T. In Vivo Imaging of Disease-Related Mitochondrial Dynamics in a Vertebrate Model System. Journal of Neuroscience 2012, 32, 16203–16212.

14. Xu, Y.; Chen, M.; Hu, B.; Huang, R.; Hu, B. In vivo Imaging of Mitochondrial Transport in Single-Axon Regeneration of Zebrafish Mauthner Cells. Frontiers in Cellular Neuroscience 2017, 11, 4.

15. Mattedi, F.; Vagnoni, A. Temporal Control of Axonal Transport: The Extreme Case of Organismal Ageing. Frontiers in Cellular Neuroscience 2019, 13, 393.

16. Misgeld, T.; Kerschensteiner, M.; Bareyre, F. M.; Burgess, R. W.; Lichtman, J. W. Imaging axonal transport of mitochondria in vivo. Nature Methods 2007-07, 4, 559–561.

17. Tosolini, A. P.; Villarroel-Campos, D.; Schiavo, G.; Sleigh, J. N. Expanding the Toolkit for jem¿In Vivo¡/em¿ Imaging of Axonal Transport. Journal of Visualized Experiments 2021,

18. Sorbara, C.; Wagner, N.; Ladwig, A.; Nikić, I.; Merkler, D.; Kleele, T.; Marinković, P.; Naumann, R.; Godinho, L.; Bareyre, F.; Bishop, D.; Misgeld, T.; Kerschensteiner, M. Pervasive Axonal Transport Deficits in Multiple Sclerosis Models. Neuron 2014, 84, 1183–1190.

19. Takihara, Y.; Inatani, M.; Eto, K.; Inoue, T.; Kreymerman, A.; Miyake, S.; Ueno, S.; Nagaya, M.; Nakanishi, A.; Iwao, K.; Takamura, Y.; Sakamoto, H.; Satoh, K.; Kondo, M.; Sakamoto, T.; Goldberg, J. L.; Nabekura, J.; Tanihara, H. In vivo imaging of axonal transport of mitochondria in the diseased and aged mammalian CNS. Proceedings of the National Academy of Sciences 2015, 112, 10515–10520.

20. Knabbe, J.; Nassal, J. P.; Verhage, M.; Kuner, T. Secretory vesicle trafficking in awake and anaesthetized mice: differential speeds in axons versus synapses. The Journal of Physiology 2018, 596, 3759–3773.

21. Chowdary, P.; Che, D.; Zhang, K.; Cui, B. Retrograde NGF Axonal Transport—Motor Coordination in the Unidirectional Motility Regime. Biophysical Journal 2015, 108, 2691–2703.

22. Haziza, S.; Mohan, N.; Loe-Mie, Y.; Lepagnol-Bestel, A.-M.; Massou, S.; Adam, M.-P.; Le, X. L.; Viard, J.; Plancon, C.; Daudin, R.; Koebel, P.; Dorard, E.; Rose, C.; Hsieh, F.-J.; Wu, C.-C.; Potier, B.; Herault, Y.; Sala, C.; Corvin, A.; Allinquant, B. et al. Fluorescent nanodiamond tracking reveals intraneuronal transport abnormalities induced by brain-disease-related genetic risk factors. Nature nanotechnology 2016, 12, 322–328.

23. Hendricks, A. G.; Perlson, E.; Ross, J. L.; Schroeder, H. W.; Tokito, M.; Holzbaur, E. L. Motor Coordination via a Tug-of-War Mechanism Drives Bidirectional Vesicle Transport. Current Biology 2010, 20, 697–702.

24. Cui, B.; Wu, C.; Chen, L.; Ramirez, A.; Bearer, E. L.; Li, W.-P.; Mobley, W. C.; Chu, S. One at a time, live tracking of NGF axonal transport using quantum dots. Proceedings of the National Academy of Sciences 2007, 104, 13666–13671.

25. Malkinson, G.; Mahou, P.; Chaudan, E.; Gacoin, T.; Sonay, A. Y.; Pantazis, P.; Beaurepaire, E.; Supatto, W. Fast In Vivo Imaging of SHG Nanoprobes with Multiphoton Light-Sheet Microscopy. ACS Photonics 2020, 7, 1036–1049.

26. Mayer, L.; Slablab, A.; Dantelle, G.; Jacques, V.; Lepagnol-Bestel, A.-M.; Perruchas, S.; Spinicelli, P.; Thomas, A.; Chauvat, D.; Simonneau, M.; Gacoin, T.; Roch, J.-F. Single KTP nanocrystals as second-harmonic generation biolabels in cortical neurons. Nanoscale 2013, 5, 8466–8471.

27. Xuan, L. L.; Zhou, C.; Slablab, A.; Chauvat, D.; Tard, C.; Perruchas, S.; Gacoin, T.; Villeval, P.; Roch, J. Photostable Second-Harmonic Generation from a Single KTiOPO4 Nanocrystal for Nonlinear Microscopy. Small 2008-09, 4, 1332–1336.

28. Gu, Y.; Sun, W.; Wang, G.; Jeftinija, K.; Jeftinija, S.; Fang, N. Rotational dynamics of cargos at pauses during axonal transport. Nature Communications 2012, 3, 1030.

29. Kaplan, L.; Ierokomos, A.; Chowdary, P.; Bryant, Z.; Cui, B. Rotation of endosomes demonstrates coordination of molecular motors during axonal transport. Science Advances 2018, 4, e1602170.

30. Donato, V. D.; Santis, F. D.; Albadri, S.; Auer, T. O.; Duroure, K.; Charpentier, M.; Concordet, J.-P.; Gebhardt, C.; Bene, F. D. An Attractive Reelin Gradient Establishes Synaptic Lamination in the Vertebrate Visual System. Neuron 2018, 97, 1049–1062.e6.

31. Insinna, C.; Baye, L. M.; Amsterdam, A.; Besharse, J. C.; Link, B. A. Analysis of a zebrafish dync1h1 mutant reveals multiple functions for cytoplasmic dynein 1 during retinal photoreceptor development. Neural Development 2010, 5, 12–12.

32. Auer, T. O.; Xiao, T.; Bercier, V.; Gebhardt, C.; Duroure, K.; Concordet, J.-P.; Wyart, C.; Suster, M.; Kawakami, K.; Wittbrodt, J.; Baier, H.; Bene, F. D. Deletion of a kinesin I motor unmasks a mechanism of homeostatic branching control by neurotrophin-3. eLife 2015, 4, e05061.

33. Steinman, J. B.; Santarossa, C. C.; Miller, R. M.; Yu, L. S.; Serpinskaya, A. S.; Furukawa, H.; Morimoto, S.; Tanaka, Y.; Nishitani, M.; Asano, M.; Zalyte, R.; On-drus, A. E.; Johnson, A. G.; Ye, F.; Nachury, M. V.; Fukase, Y.; Aso, K.; Foley, M. A.; Gelfand, V. I.; Chen, J. K. et al. Chemical structure-guided design of dynapyrazoles, cell-permeable dynein inhibitors with a unique mode of action. eLife 2017, 6, e25174.

34. Niell, C. M.; Smith, S. J. Functional Imaging Reveals Rapid Development of Visual Response Properties in the Zebrafish Tectum. Neuron 2005, 45, 941–951.

35. Eichel, K.; Uenaka, T.; Belapurkar, V.; Lu, R.; Cheng, S.; Pak, J. S.; Taylor, C. A.; Südhof, T. C.; Malenka, R.; Wernig, M.; Ozkan, E.; Perrais, D.; Shen, K. Endocytosis in the axon initial segment maintains neuronal polarity. Nature 2022, 609, 128–135.

36. Grimaud, B.; Terras, F.; Marquier, F. biophotlumin/mint: v0.1.4 (v0.1.4). 2022; https://doi.org/10.5281/zenodo.6669513.

37. Chou, Q.-L.; Alik, A.; Marquier, F.; Melki, R.; Treussart, F.; Simonneau, M. Impact of α-Synuclein Fibrillar Strains and β-Amyloid Assemblies on Mouse Cortical Neurons Endo-Lysosomal Logistics. eNeuro 2022, 9, ENEURO.0227-21.2022.

38. Boyd, R. W. Nonlinear Optics, 4th ed.; Elsevier Science, 2020.

39. Kusumi, A.; Sako, Y.; Yamamoto, M. Confined lateral diffusion of membrane receptors as studied by single particle tracking (nanovid microscopy). Effects of calcium-induced differentiation in cultured epithelial cells. Biophysical Journal 1993, 65, 2021–2040.

40. Bercier, V.; Rosello, M.; Bene, F. D.; Revenu, C. Zebrafish as a Model for the Study of Live in vivo Processive Transport in Neurons. Frontiers in Cell and Developmental Biology 2019, 7, 17.

41. Rudin, L. I.; Osher, S.; Fatemi, E. Nonlinear total variation based noise removal algorithms. Physica D: Nonlinear Phenomena 1992, 60, 259–268.

42. Redwine, W. B.; DeSantis, M. E.; Hollyer, I.; Htet, Z. M.; Tran, P. T.; Swanson, S. K.; Florens, L.; Washburn, M. P.; Reck-Peterson, S. L. The human cytoplasmic dynein interactome reveals novel activators of motility. eLife 2017, 6, e28257.

43. Martin, M.; Iyadurai, S. J.; Gassman, A.; Gindhart, J. G.; Hays, T. S.; Saxton, W. M. Cytoplasmic Dynein, the Dynactin Complex, and Kinesin Are Interdependent and Essential for Fast Axonal Transport. Molecular Biology of the Cell 1999, 10, 3717–3728.

44. Deacon, S. W.; Serpinskaya, A. S.; Vaughan, P. S.; Fanarraga, M. L.; Vernos, I.; Vaughan, K. T.; Gelfand, V. I. Dynactin is required for bidirectional organelle transport. The Journal of Cell Biology 2003, 160, 297–301.

45. Herbert, A. L.; Fu, M.-m.; Drerup, C. M.; Gray, R. S.; Harty, B. L.; Ackerman, S. D.; O’Reilly-Pol, T.; Johnson, S. L.; Nechiporuk, A. V.; Barres, B. A.; Monk, K. R. Dynein/dynactin is necessary for anterograde transport of Mbp mRNA in oligodendrocytes and for myelination in vivo. Proceedings of the National Academy of Sciences of the United States of America 2017, 114, E9153–E9162.

46. Encalada, S. E.; Szpankowski, L.; Xia, C.-h.; Goldstein, L. S. Stable Kinesin and Dynein Assemblies Drive the Axonal Transport of Mammalian Prion Protein Vesicles. Cell 2011-02, 144, 551–565.

47. Konjikusic, M. J.; Gray, R. S.; Wallingford, J. B. The developmental biology of kinesins. Developmental Biology 2020, 469, 26–36.

48. Campbell, P. D.; Marlow, F. L. Temporal and tissue specific gene expression patterns of the zebrafish kinesin-1 heavy chain family, kif5s, during development. Gene Expression Patterns 2013-10, 13, 271–279.

49. Zhao, C.; Omori, Y.; Brodowska, K.; Kovach, P.; Malicki, J. Kinesin-2 family in vertebrate ciliogenesis. Proceedings of the National Academy of Sciences 2012, 109, 2388–2393.

50. Takeda, S.; Yamazaki, H.; Seog, D.-H.; Kanai, Y.; Terada, S.; Hirokawa, N. Kinesin Superfamily Protein 3 (Kif3) Motor Transports Fodrin-Associating Vesicles Important for Neurite Building. The Journal of Cell Biology 2000, 148, 1255–1266.

51. Schroeder, H.; Hendricks, A. G.; Ikeda, K.; Shuman, H.; Rodionov, V.; Ikebe, M.; Goldman, Y.; Holzbaur, E. Force-Dependent Detachment of Kinesin-2 Biases Track Switching at Cytoskeletal Filament Intersections. Biophysical Journal 2012, 103, 48–58.

52. White, R. M.; Sessa, A.; Burke, C.; Bowman, T.; LeBlanc, J.; Ceol, C.; Bourque, C.; Dovey, M.; Goessling, W.; Burns, C. E.; Zon, L. I. Transparent Adult Zebrafish as a Tool for In Vivo Transplantation Analysis. Cell Stem Cell 2008, 2.

53. Ge, R.; Zhou, Y.; Peng, R.; Wang, R.; Li, M.; Zhang, Y.; Zheng, C.; Wang, C. Conservation of the STING-Mediated Cytosolic DNA Sensing Pathway in Zebrafish. Journal of Virology 2015, 89, 7696–7706.

54. Passoni, G.; Langevin, C.; Palha, N.; Mounce, B. C.; Briolat, V.; Affaticati, P.; Job, E. D.; Joly, J.-S.; Vignuzzi, M.; Saleh, M.-C.; Herbomel, P.; Boudinot, P.; Levraud, J.-P. Imaging of viral neuroinvasion in the zebrafish reveals that Sindbis and chikungunya viruses favour different entry routes. Disease Models & Mechanisms 2017, 10, 847–857.

55. Guerra-Varela, J.; Baz-Martínez, M.; Silva-Álvarez, S. D.; Losada, A. P.; Quiroga, M. I.; Collado, M.; Rivas, C.; Sánchez, L. Susceptibility of Zebrafish to Vesicular Stomatitis Virus Infection. Zebrafish 2018, 15, 124–132.

56. Bhattarai, P.; Thomas, A.; Cosacak, M.; Papadimitriou, C.; Mashkaryan, V.; Froc, C.; Reinhardt, S.; Kurth, T.; Dahl, A.; Zhang, Y.; Kizil, C. IL4/STAT6 Signaling Activates Neural Stem Cell Proliferation and Neurogenesis upon Amyloid-β42 Aggregation in Adult Zebrafish Brain. Cell Reports 2016, 17, 941–948.

57. Dukes, A. A.; Bai, Q.; Laar, V. S. V.; Zhou, Y.; Ilin, V.; David, C. N.; Agim, Z. S.; Bonkowsky, J. L.; Cannon, J. R.; Watkins, S. C.; Croix, C. M. S.; Burton, E. A.; Berman, S. B. Live imaging of mitochondrial dynamics in CNS dopaminergic neurons in vivo demonstrates early reversal of mitochondrial transport following MPP+ exposure. Neurobiology of Disease 2016, 95, 238–249.

58. Kim, G.-H. J.; Mo, H.; Liu, H.; Wu, Z.; Chen, S.; Zheng, J.; Zhao, X.; Nucum, D.; Shortland, J.; Peng, L.; Elepano, M.; Tang, B.; Olson, S.; Paras, N.; Li, H.; Renslo, A. R.; Arkin, M. R.; Huang, B.; Lu, B.; Sirota, M. et al. A zebrafish screen reveals Renin-angiotensin system inhibitors as neuroprotective via mitochondrial restoration in dopamine neurons. eLife 2021, 10, e69795.

59. Medeiros, G. d.; Kromm, D.; Balazs, B.; Norlin, N.; Günther, S.; Izquierdo, E.; Ronchi, P.; Komoto, S.; Krzic, U.; Schwab, Y.; Peri, F.; Renzis, S. d.; Leptin, M.; Rauzi, M.; Hufnagel, L. Cell and tissue manipulation with ultrashort infrared laser pulses in light-sheet microscopy. Scientific Reports 2020, 10, 1942.

60. Yang, B.; Treweek, J.; Kulkarni, R.; Deverman, B.; Chen, C.-K.; Lubeck, E.; Shah, S.; Cai, L.; Gradinaru, V. Single-Cell Phenotyping within Transparent Intact Tissue through Whole-Body Clearing. Cell 2014, 158.

61. Bestvater, F.; Spiess, E.; Stobrawa, G.; Hacker, M.; Feurer, T.; Porwol, T.; Berchner-Pfannschmidt, U.; Wotzlaw, C.; Acker, H. Two-photon fluorescence absorption and emission spectra of dyes relevant for cell imaging. Journal of Microscopy 2002, 208, 108–115.

62. Linkert, M.; Rueden, C. T.; Allan, C.; Burel, J.-M.; Moore, W.; Patterson, A.; Loranger, B.; Moore, J.; Neves, C.; MacDonald, D.; Tarkowska, A.; Sticco, C.; Hill, E.; Rossner, M.; Eliceiri, K. W.; Swedlow, J. R. Metadata matters: access to image data in the real world. The Journal of Cell Biology 2010, 189.

63. Allan, D. B.; Caswell, T.; Keim, N. C.; van der Wel, C. M.; Verweij, R. W. soft-matter/trackpy: Trackpy v0.5.0. 2021; https://doi.org/10.5281/zenodo.4682814.

64. Crocker, J. C.; Grier, D. G. Methods of Digital Video Microscopy for Colloidal Studies. Journal of Colloid and Interface Science 1996, 179, 298–310.

65. Diamond, S.; Boyd, S. CVXPY: A Python-embedded modeling language for convex optimization. Journal of Machine Learning Research 2016, 17, 1–5.

66. Agrawal, A.; Verschueren, R.; Diamond, S.; Boyd, S. A rewriting system for convex optimization problems. Journal of Control and Decision 2018, 5, 42–60.

67. O’Donoghue, B.; Chu, E.; Parikh, N.; Boyd, S. SCS: Splitting Conic Solver, version 3.2.1. https://github.com/cvxgrp/scs, 2021.

68. O’Donoghue, B.; Chu, E.; Parikh, N.; Boyd, S. Conic Optimization via Operator Splitting and Homogeneous Self-Dual Embedding. Journal of Optimization Theory and Applications 2016, 169, 1042–1068.

## References

1. Balzarotti, F.; Eilers, Y.; Gwosch, K. C.; Gynnå, A. H.; Westphal, V.; Stefani, F. D.; Elf, J.; Hell, S. W. Nanometer resolution imaging and tracking of fluorescent molecules with minimal photon fluxes. Science 2017, 355, 606–612.

2. Masullo, L. A.; Lopez, L. F.; Stefani, F. D. A common framework for single-molecule localization using sequential structured illumination. Biophysical Reports 2022, 2, 100036.

3. Deschout, H.; Zanacchi, F. C.; Mlodzianoski, M.; Diaspro, A.; Bewersdorf, J.; Hess, S. T.; Braeckmans, K. Precisely and accurately localizing single emitters in fluorescence microscopy. Nature Methods 2014, 11, 253–266.

